# Mitochondrial Transport in Symmetric and Asymmetric Axons with Multiple Branching Junctions: A Computational Study

**DOI:** 10.1101/2023.02.22.529604

**Authors:** Ivan A. Kuznetsov, Andrey V. Kuznetsov

## Abstract

We explore the impact of multiple branching junctions in axons on the mean age of mitochondria and their age density distributions in demand sites. The study looked at mitochondrial concentration, mean age, and age density distribution in relation to the distance from the soma. We developed models for a symmetric axon containing 14 demand sites and an asymmetric axon containing 10 demand sites. We examined how the concentration of mitochondria changes when an axon splits into two branches at the branching junction. We also studied whether mitochondria concentrations in the branches are affected by what proportion of mitochondrial flux enters the upper branch and what proportion of flux enters the lower branch. Additionally, we explored whether the distributions of mitochondria mean age and age density in branching axons are affected by how the mitochondrial flux splits at the branching junction. When the mitochondrial flux is split unevenly at the branching junction of an asymmetric axon, with a greater proportion of the flux entering the longer branch, the average age of mitochondria (system age) in the axon increases. Our findings elucidate the effects of axonal branching on mitochondria age. Mitochondria aging is the focus of this study as recent research suggests it may be involved in neurodegenerative disorders, such as Parkinson’s disease.

## 1. Introduction

Dopaminergic (DA) neurons in the substantia nigra pars compacta die during Parkinson’s disease (PD) (Chinta and Andersen 2005; Guo et al. 2018; Xu et al. 2017). The axonal arbors of these neurons are large and unmyelinated, and the morphology of their axons is very complex with multiple branching junctions. In humans, a single DA axon can form more than a million synapses (Matsuda et al. 2009). Maintaining such a large arbor requires a lot of energy, and energy supply problems may be a possible explanation for the death of DA neurons in PD (Bolam and Pissadaki 2012). This motivates our interest in axons that have multiple branching junctions.

Mitochondria are important for regulating physiology and cellular functions of neurons (Pekkurnaz and Wang 2022). Continuous transport of mitochondria in axons (Mogre et al. 2020) is critical for maintaining mitochondrial health. To maintain good mitochondrial health in an axon, it is necessary to periodically replace either a complete mitochondrion or the proteins within it (Misgeld and Schwarz 2017).

Our specific interest is mitochondria traffic in axons with multiple branches and the effect of branched paths on mitochondrial aging (Vanhauwaert et al. 2019). Due to the large size of mitochondria, their diffusivity is negligible (Trushina 2016), and mitochondria transport is driven by molecular motors (Puttrich et al. 2022). Anterograde transport of mitochondria is driven by kinesin-1 while retrograde transport is driven by cytoplasmic dynein (Melkov and Abdu 2018; Kruppa and Buss 2021). Mitochondria can become anchored at various energy demand sites in the axon (Hollenbeck and Saxton 2005; Lees et al. 2020; Lewis et al. 2016).

Impaired mitochondrial transport in neurons plays a significant role in neurodegeneration (Sheng and Cai 2012; Correia et al. 2016). CNS regeneration can be promoted by turning off anchoring of mitochondria (Cheng et al. 2022). Transport of new mitochondria is essential for axon regeneration following injury (McElroy et al. 2023). We are therefore interested in the distributions of mitochondria concentration and mitochondria age in branching axons, and how these distributions are impacted by the axon geometry. We investigate how the concentration of mitochondria changes when an axon splits into two branches and how axon symmetry/asymmetry affects this concentration. We are also interested in the distribution of mitochondria mean age and age density when an axon splits into branches. Finally, we are interested in how the symmetry or asymmetry in the splitting of mitochondrial flux at the branching junction affects the mean age and age density distribution of mitochondria in the branches and the axon segment before the junction. The obtained results may help in understanding the relationship between the asymmetry in axon morphology and mitochondrial aging, which is important because mitochondrial aging may contribute to the development of neurodegenerative conditions.

Karamched and Bressloff (2017) used mean-field equations to investigate transport of motor-cargo complexes to axons with multiple branches. Patel et al. (2013) developed equations for mitochondrial health, a quantity used as a proxy for mitochondrial membrane potential. Agrawal and Koslover (2021) used the mean-field theory to create models that simulated mitochondrial health as a decaying component. They applied their models to investigating the health of mitochondria in both straight and branched axons.

Kuznetsov and Kuznetsov (2022a, 2022b, 2023) developed an alternative model of mitochondria transport in an axon that is based on simulating mitochondria transport and accumulation in various energy demand sites. The motivation was to apply powerful methods developed for compartmental systems (Rasmussen et al. 2016; Metzler et al. 2018; Metzler and Sierra 2018; Chappelle et al. 2023) to determine mean age and age density distributions of mitochondria in demand sites located at different distances from the soma. The model developed in our research was obtained by extending the model of mitochondria transport suggested in Kuznetsov and Kuznetsov (2022a, 2022b, 2023) to symmetric and asymmetric axons with multiple branching junctions (Fig. 1).

**Fig. 1.**
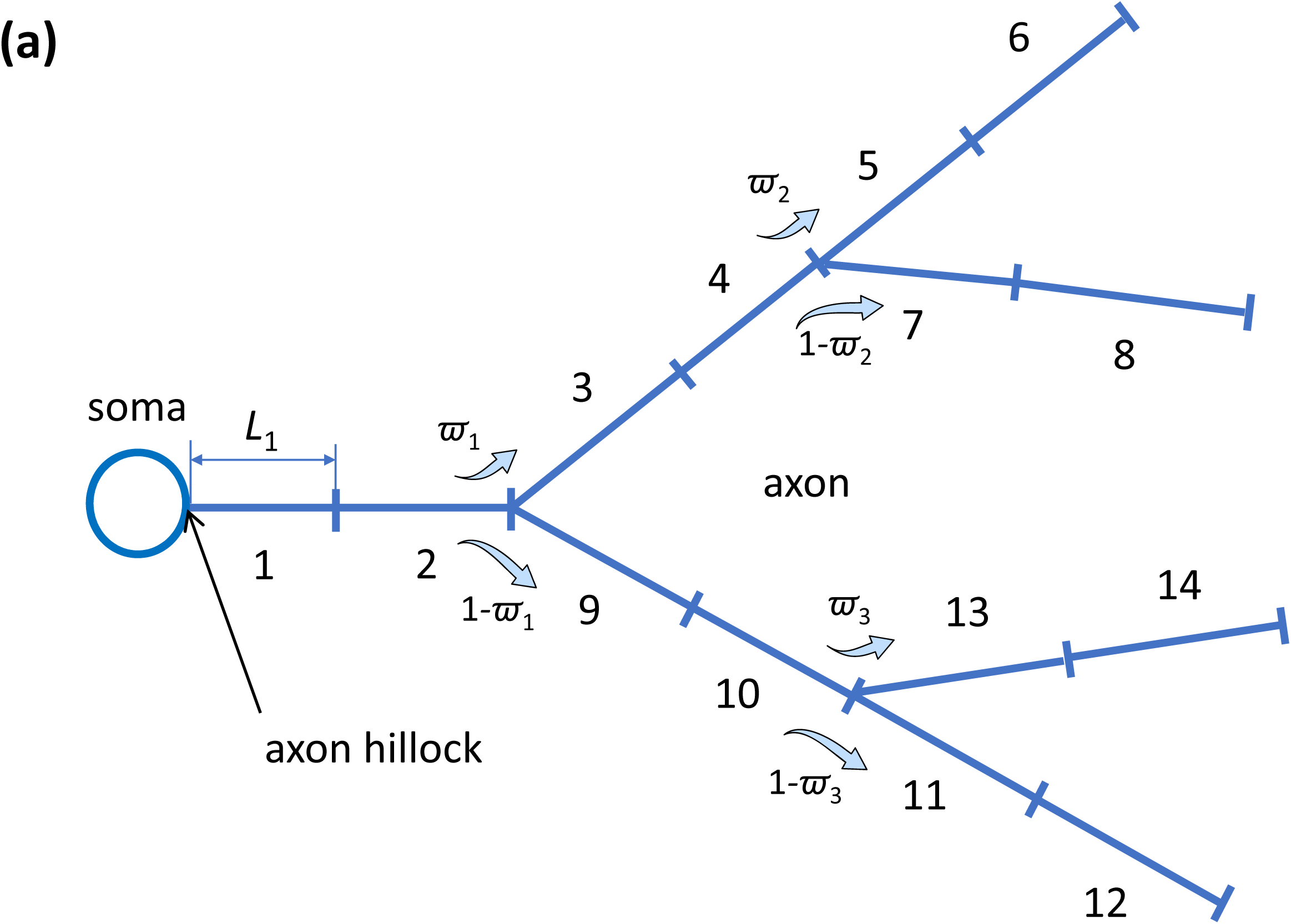
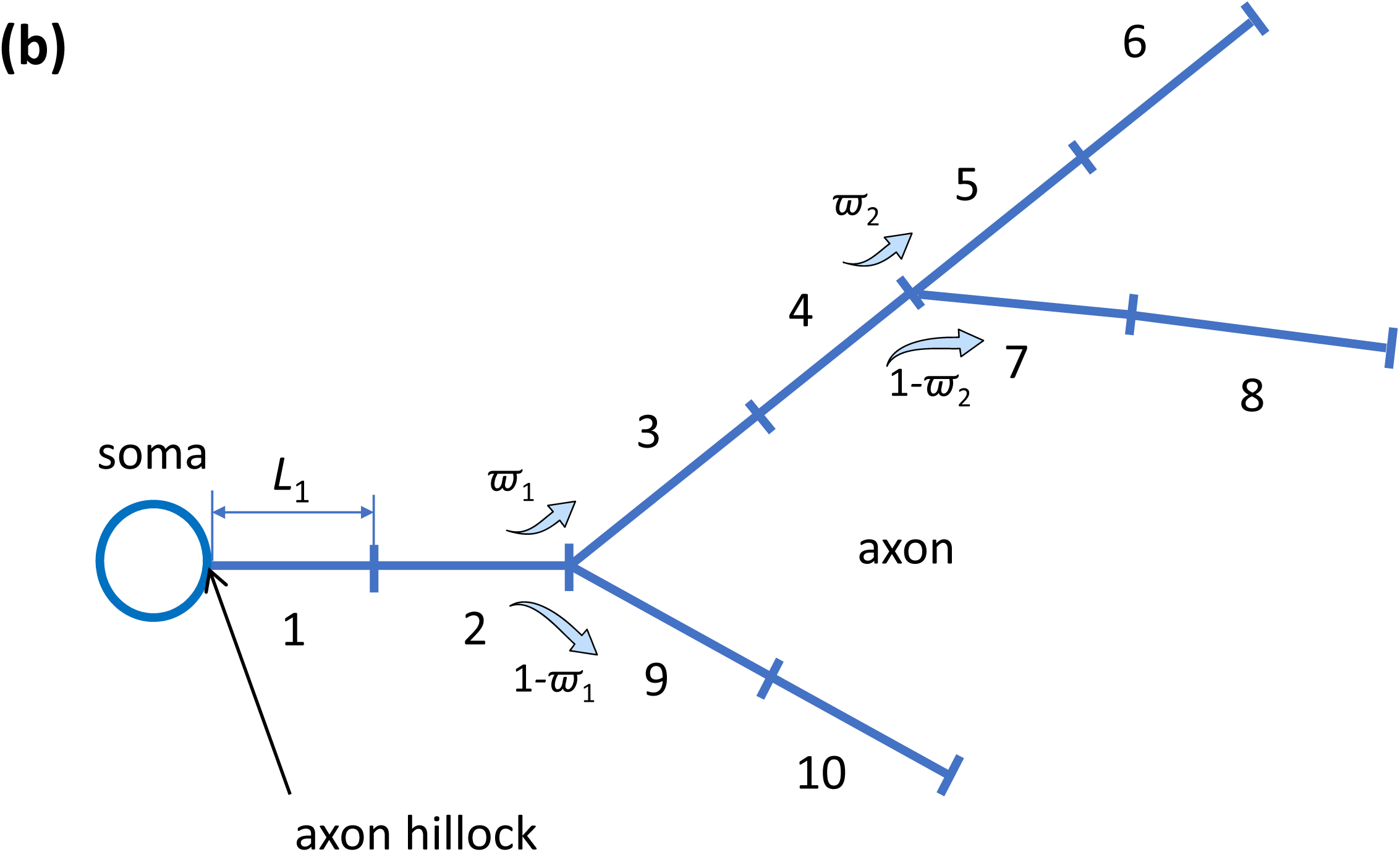
A schematic representation of a neuron with a symmetric branched axon (a) and an asymmetric branched axon (b). The diagram shows the compartment length closest to the proximal demand site. We assumed that all other compartments have equal lengths. The figure was created with the assistance of Servier Medical Art and is licensed under a Creative Commons Attribution 3.0 Generic License, available at http://smart.servier.com.

## 2. Materials and models

The transport of mitochondria in the axon is modeled using the multi-compartment model (Anderson 1983; Jacquez 1985; Jacquez and Simon 1993). This model was selected because we desired to simulate the mean ages and distributions of age density for mitochondria in various parts of the axon and its branches. To accomplish this, we have applied the techniques developed in Metzler et al. (2018) and Metzler and Sierra (2018), enabling particle age calculation in compartmental systems.

Intending to explore the impact of unequal length of axon branches on the age distribution of mitochondria in the axon, we represented an axon with symmetric branches by 14 demand sites (Fig. 1a) and an axon with asymmetric branches by 10 demand sites (Fig. 1b). Mitochondria exhibit anterograde and retrograde transport and also frequently stop for a short time at various locations during transport. Additionally, they display changes in direction, but each mitochondrion has a strong preference for traveling in a particular direction for longer distances (Saxton and Hollenbeck 2012; Alsina et al. 2017). To simulate these movements and stops in our model, we divided each demand site into three separate compartments: one for mitochondria moving in the anterograde direction, one for stationary mitochondria, and one for mitochondria moving in the retrograde direction mitochondria (as illustrated in Fig. 2a,b). The mitochondria are able to transition between these compartments, and they may also move from one demand site to the next or return to the previous demand site.

**Fig. 2.**
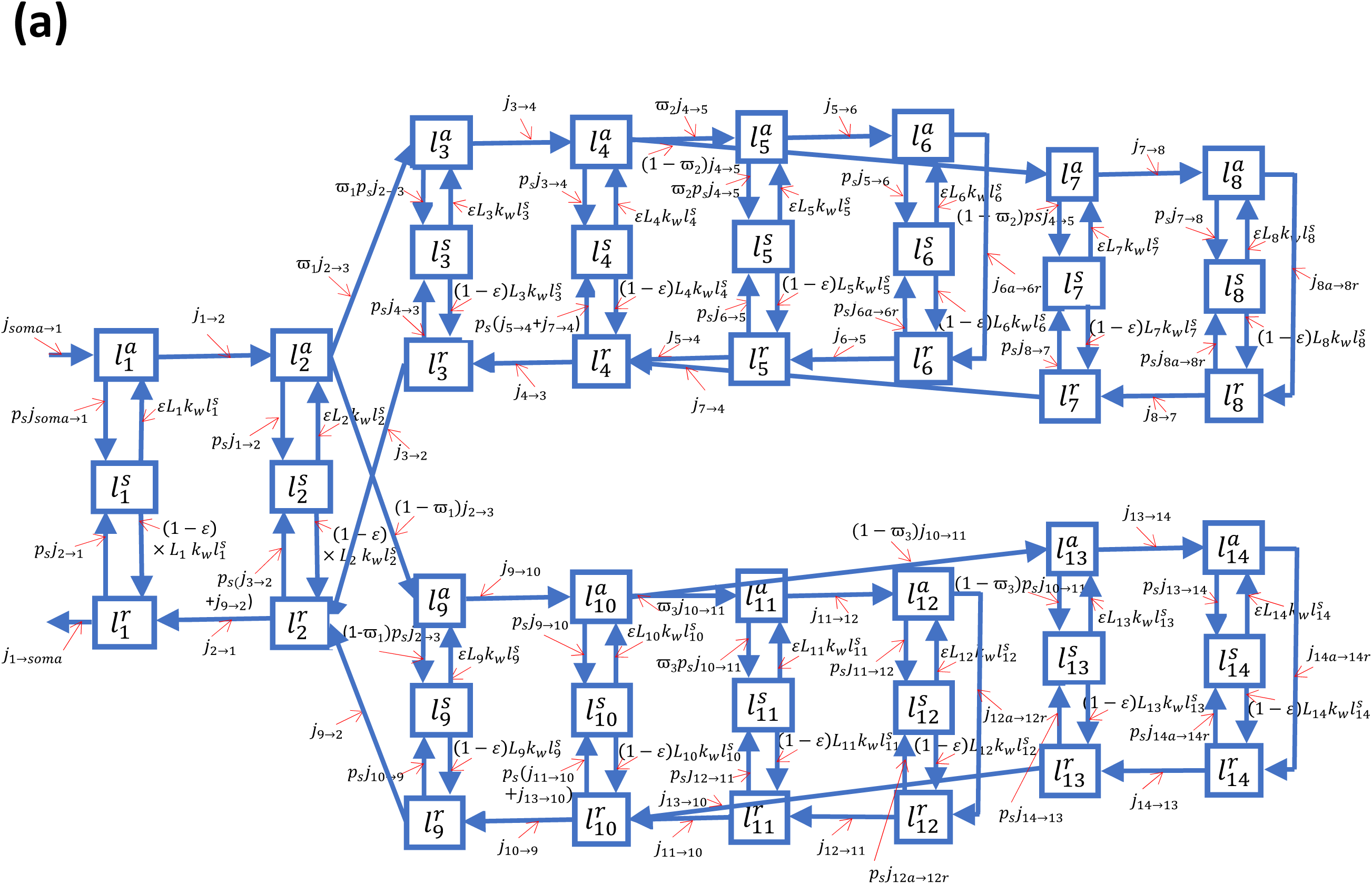
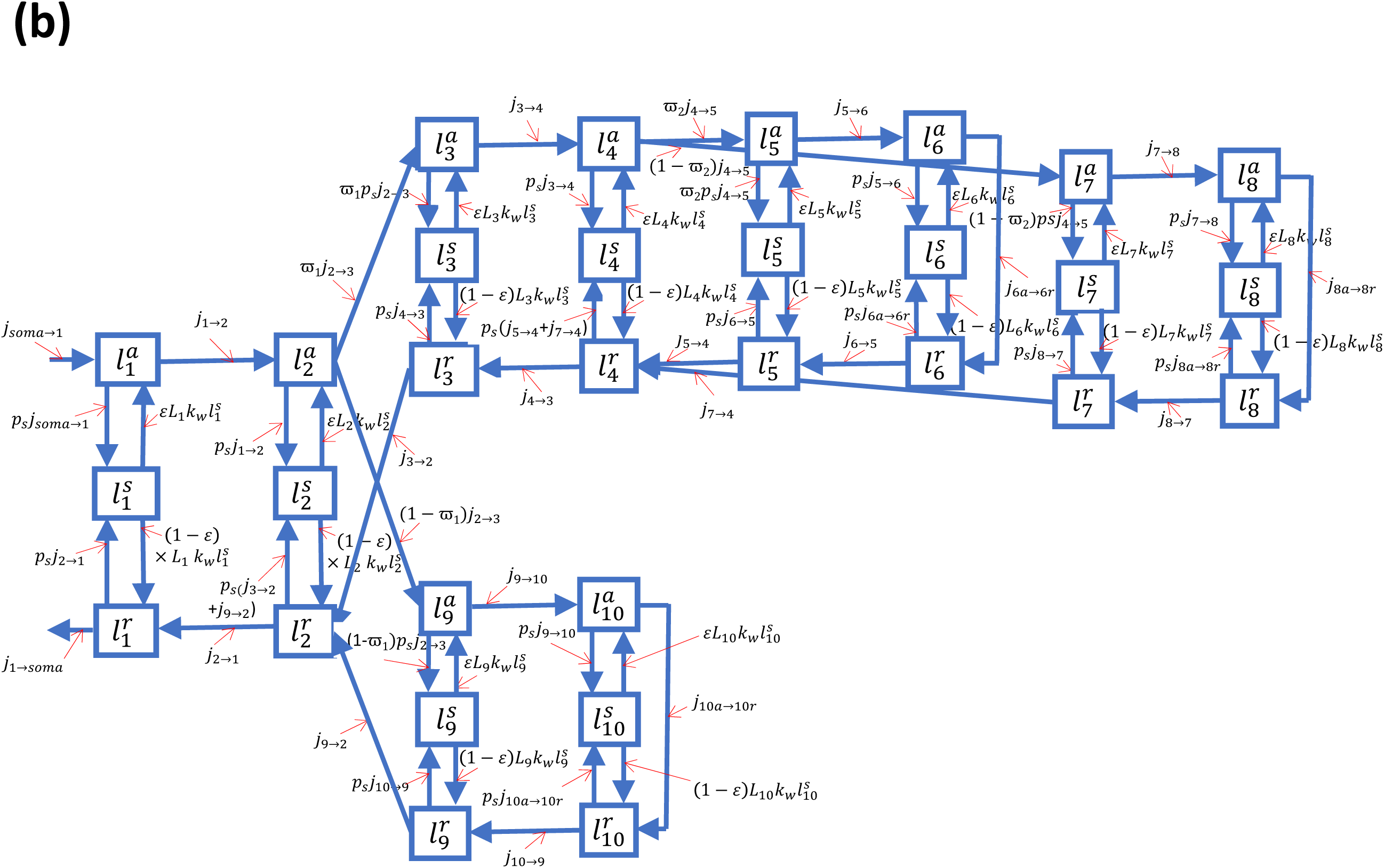
A kinetic diagram representing a compartmental model of a branched axon. The compartments symbolize demand sites in the terminal, each represented by three compartments containing anterograde, stationary, and retrograde mitochondria. The diagram uses arrows to show the transport of mitochondria between transiting (anterograde and retrograde) states in neighboring demand sites. We assumed that mitochondria entering the terminal have the initial age of zero. Arrows in the diagram also depict the capture of mobile mitochondria into stationary states and their release from these states. Part (a) of the diagram displays a symmetric branched axon, while part (b) displays an asymmetric branched axon.

Mitochondria are constantly undergoing fusion and fission (Bockler et al. 2017). To track the mitochondria with varying lengths, a mean-field equation model was not suitable. Instead, we lumped the lengths of all the mitochondria present in a single compartment. The model thus simulates the total length of mitochondria within a compartment, as this is a conserved property. Since the axons are elongated in one direction, we characterized the mitochondrial concentration by the total length of mitochondria in a given kinetic state (anterograde, stationary, or retrograde) within a unit length of the axon. The unit of measurement for mitochondrial concentration thus is (μm of mitochondria)/μm.

In our model, the symmetric axon (Fig. 1a) has 42 compartments, each simulating 42 different mitochondrial concentrations, while the asymmetric axon (Fig. 1b) has 30 compartments, each simulating 30 different mitochondrial concentrations. In this compartmental system, time (*t*) is the only independent variable. The dependent variables and model parameters are summarized in Table 1 and Table 2, respectively.

**Table 1.** Dependent variables in the compartmental model of mitochondria transport in the axon.

| Symbol | Definition | Units |
| --- | --- | --- |
| $l_i^a$ | Concentration, per unit length of the axon, of anterograde mitochondria in the compartment by the $i$-th demand site | ($\mu\text{m}$ of mitochondria)/$\mu\text{m}$ |
| $l_i^s$ | Concentration, per unit length of the axon, of stationary mitochondria in the compartment by the $i$-th demand site | ($\mu\text{m}$ of mitochondria)/$\mu\text{m}$ |
| $l_i^r$ | Concentration, per unit length of the axon, of retrograde mitochondria in the compartment by the $i$-th demand site | ($\mu\text{m}$ of mitochondria)/$\mu\text{m}$ |
| $j_{a \rightarrow b}$ | Flux of mitochondria from one compartment ($a$) to another compartment ($b$), which is represented by the total length of mitochondria that are transported from compartment ($a$) to compartment ($b$) per unit of time | ($\mu\text{m}$ of mitochondria) $\text{s}^{-1}$ |

**Table 2.**
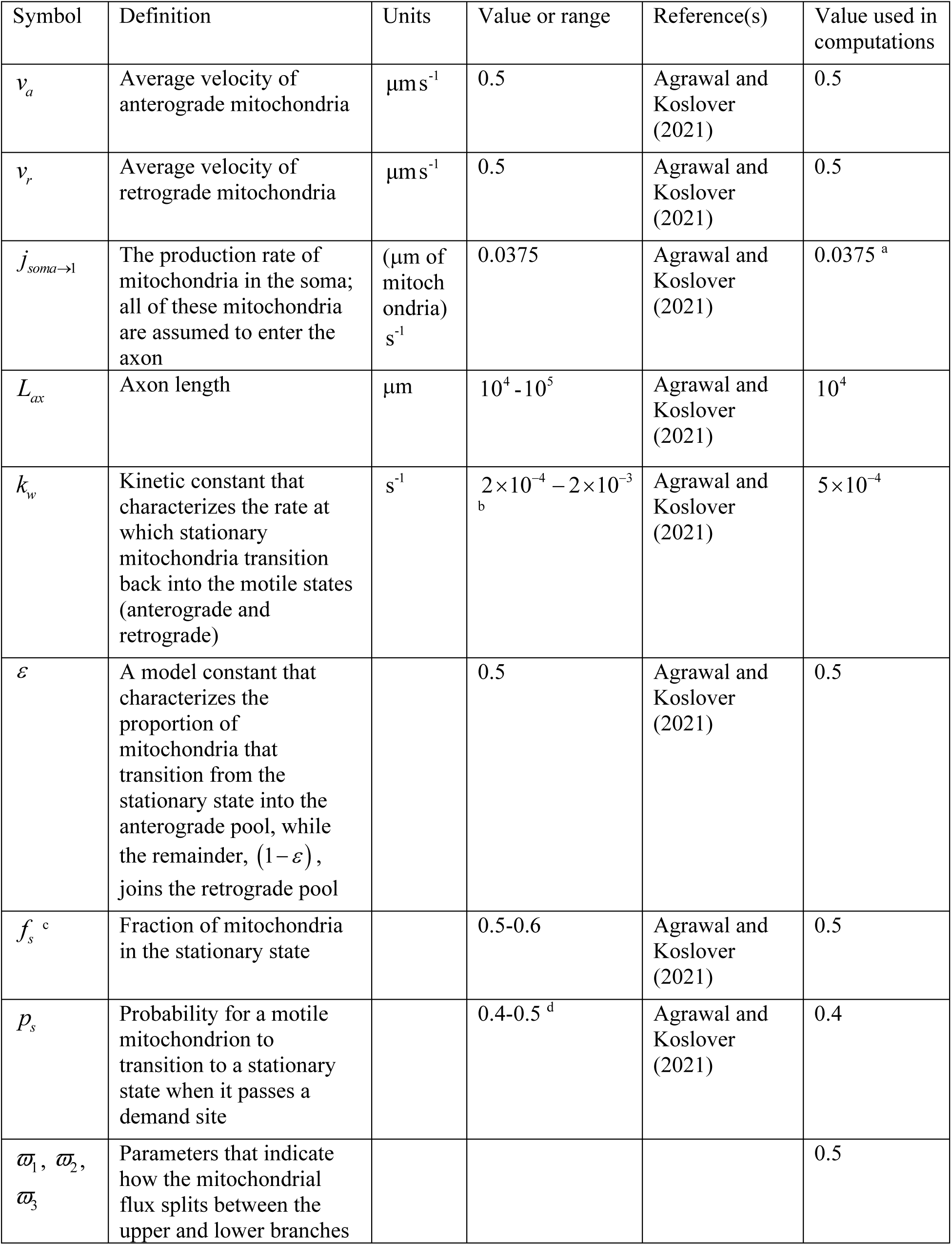
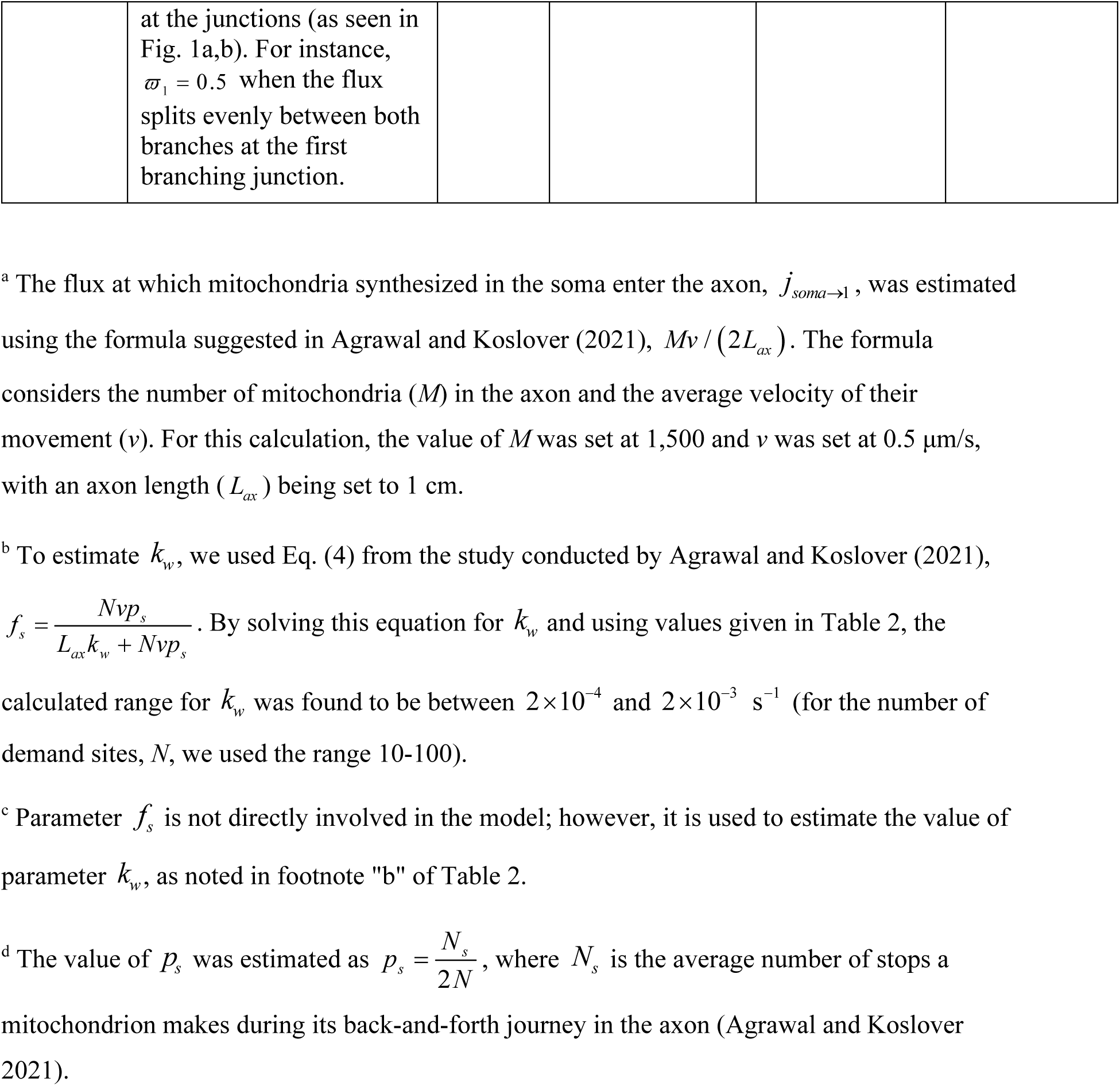
Parameters in the model of mitochondrial transport in the axon.

### 2.1. Equations that express the conservation of the total length of mitochondria in various compartments for an axon with two symmetric branches (Figs. 1a and 2a)

Eqs. (1)-(42) presented below are derived by applying the conservation of the total length of mitochondria in each of the 42 compartments illustrated in Fig. 2a. The equation that represents the conservation of the total length of stationary mitochondria in the most proximal demand site (# 1) is expressed as follows:

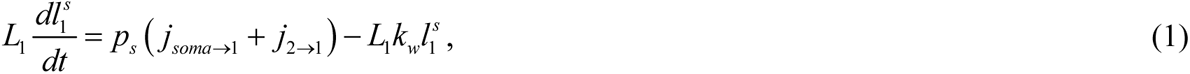

where 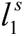 is the concentration of stationary mitochondria in the most proximal compartment and *L*_1_ is the length of the axonal compartments around the most proximal demand site. For axons shown in Fig. 1a and 1b, it was assumed that *L*_1_ = *L_ax_*/ 6. The lengths of all other axonal compartments (*L*_2_, *L*_3_, etc.) were assumed to be the same. Also, *p_s_* is the probability of a mobile mitochondrion transitioning to a stationary state as it passes a demand site, *k_w_* is the kinetic constant that characterizes the rate at which stationary mitochondria reenter the mobile states (anterograde and retrograde), and *j_soma_*_→1_ represents the rate of mitochondrial production in the soma. The fluxes between the compartments containing mobile mitochondria, *j_a_*_→_*_b_*, are given by Eqs. (43)-(70) below.

The conservation of the total length of anterograde mitochondria in the most proximal demand site (# 1) is expressed by the following equation (Fig. 2a):

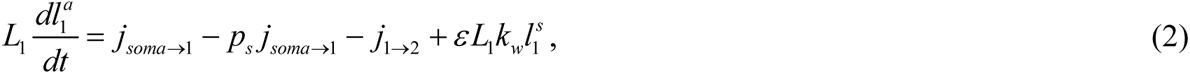

where 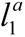 is the concentration of anterograde mitochondria in the most proximal compartment and *ε* is a model constant that simulates what proportion of mitochondria released from the stationary state will join the anterograde pool, with the remaining proportion, (1− *ε*), joining the retrograde pool.

The conservation of the total length of retrograde mitochondria in the most proximal demand site (# 1) is given by the following equation (Fig. 2a):

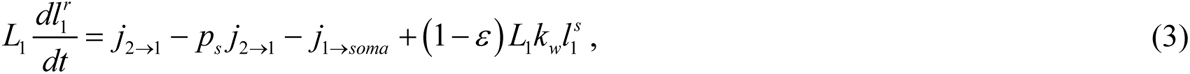

where 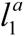 is the concentration of retrograde mitochondria in the most proximal compartment. Stating the conservation of the total length of mitochondria in the compartments that hold stationary, anterograde, and retrograde mitochondria around other demand sites (#2, …, 14) leads to the following equations:

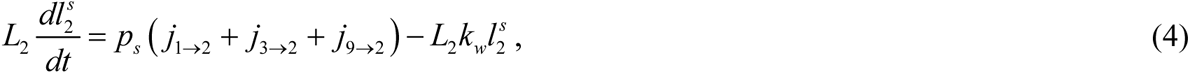

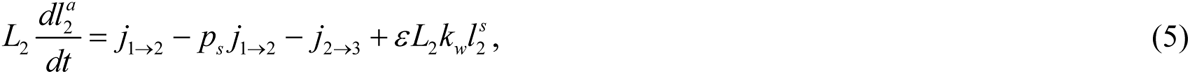

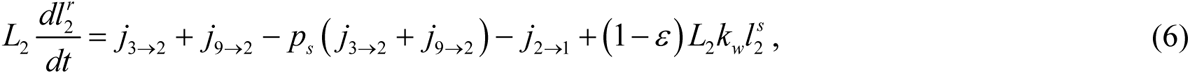

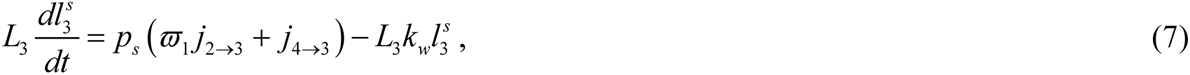

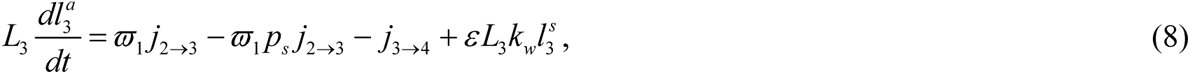

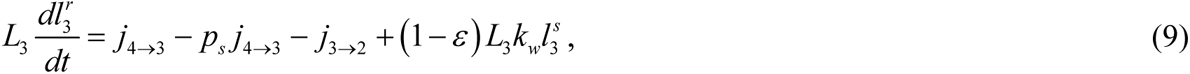

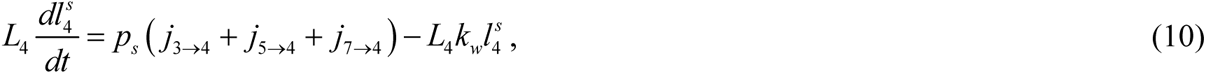

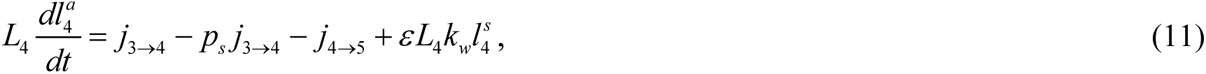

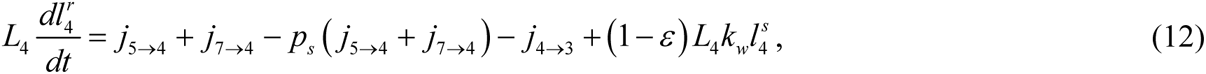

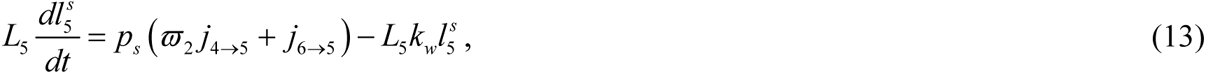

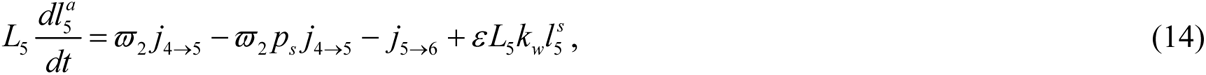

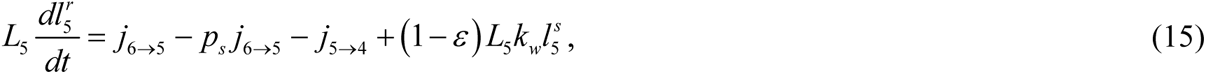

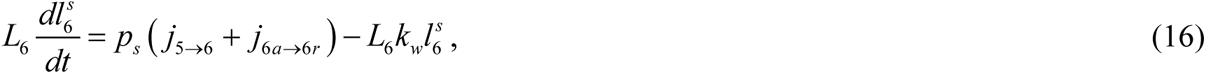

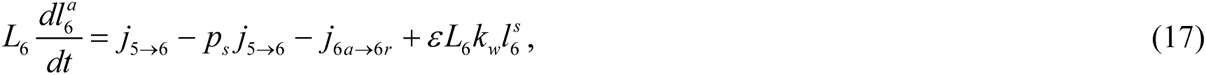

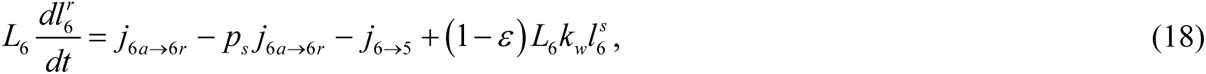

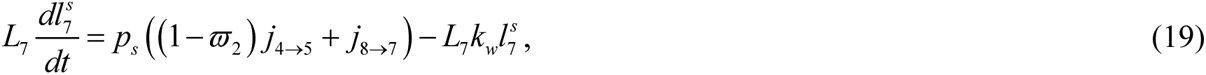

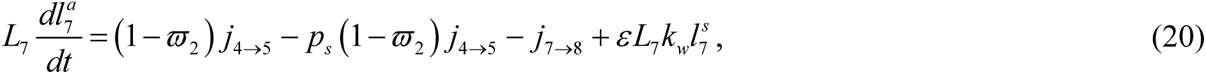

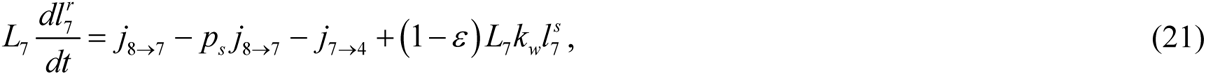

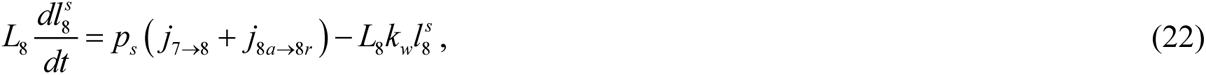

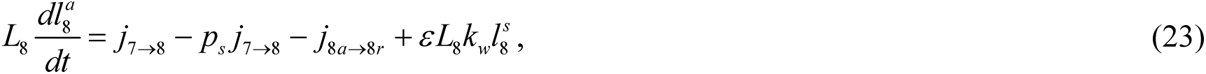

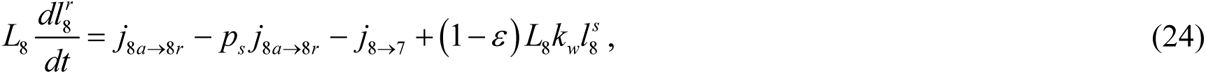

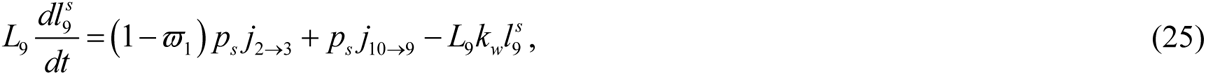

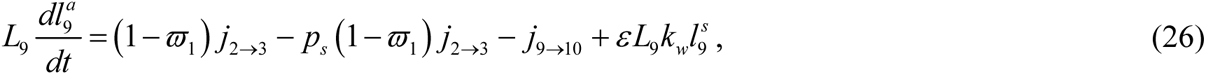

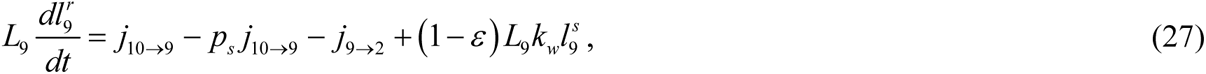

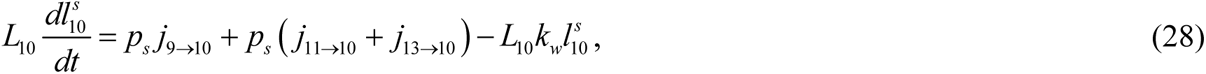

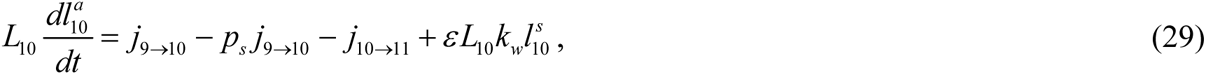

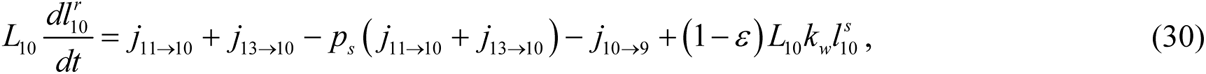

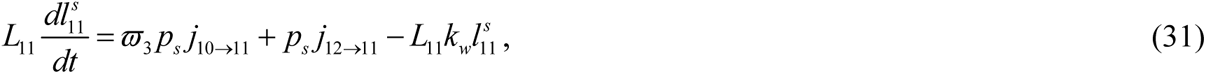

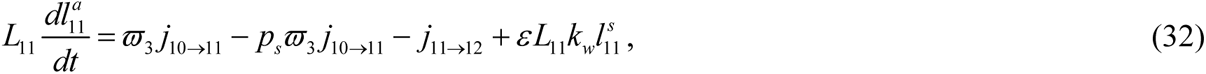

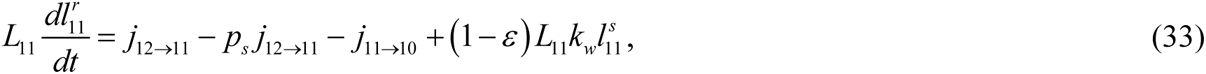

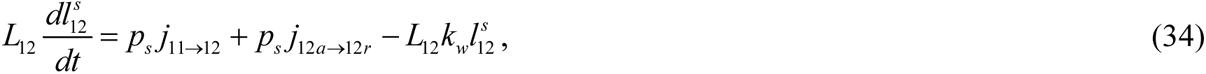

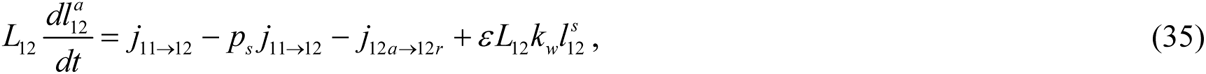

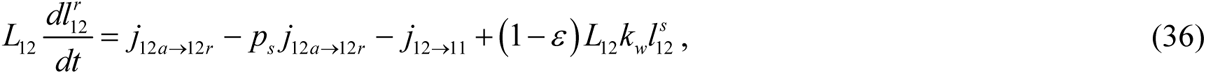

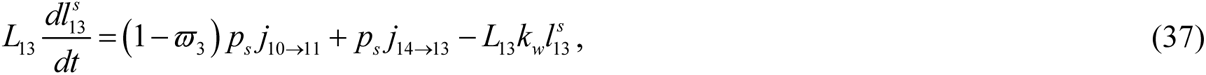

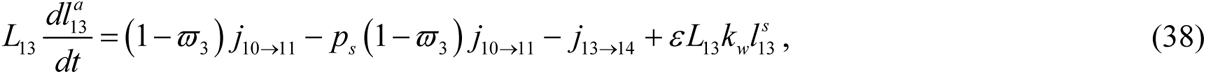

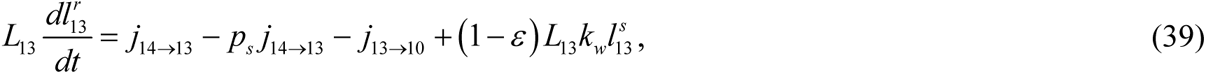

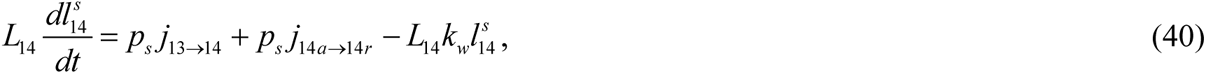

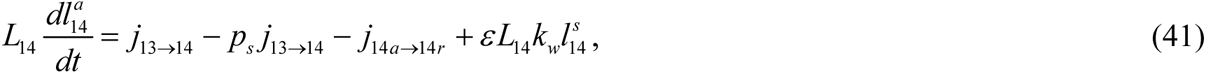

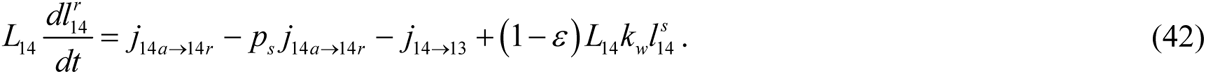

Eqs. (1)-(42) must be supplemented by equations for the mitochondrial fluxes between the demand sites. These fluxes are displayed by arrows in Fig. 2a. *j_soma_*_→1_ is an input model parameter that characterizes the rate of mitochondria production in the soma (according to our model, they all enter the axon); equations for other anterograde fluxes (Fig. 2a) are

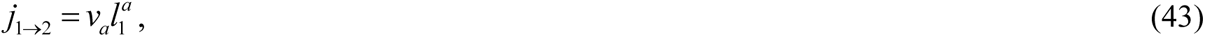

where *v_a_* is the average velocity of motile mitochondria moving anterogradely.

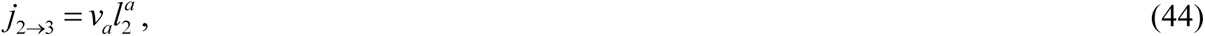

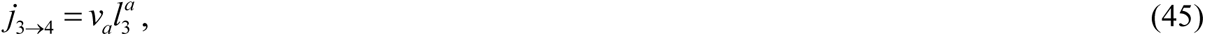

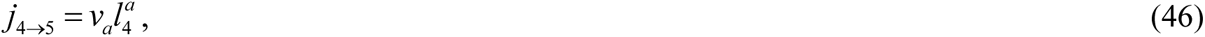

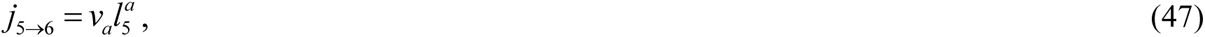

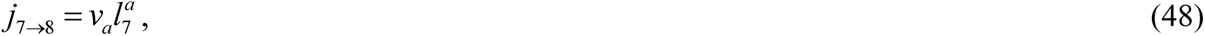

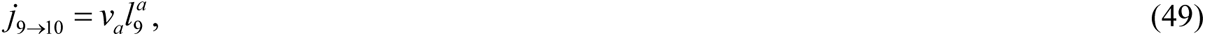

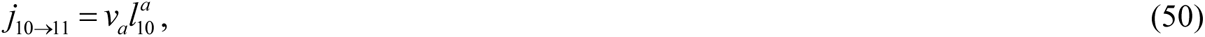

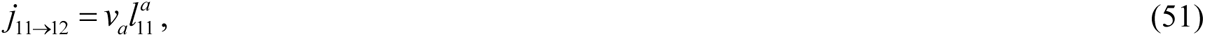

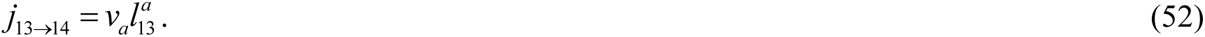

Equations for retrograde fluxes (Fig. 2a) are

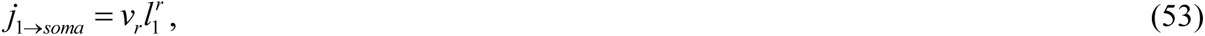

where *v_r_* is the average velocity of motile mitochondria that are moving retrogradely.

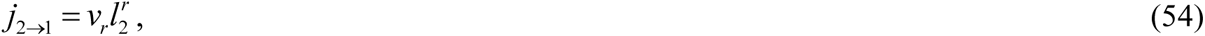

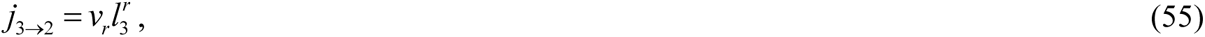

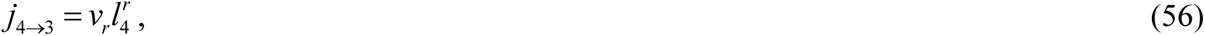

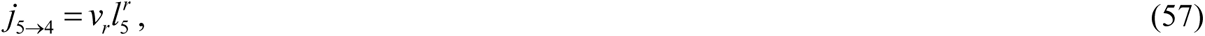

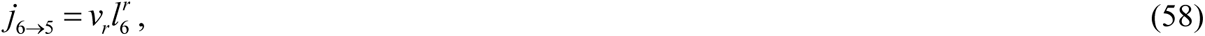

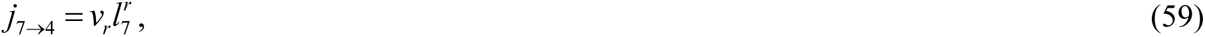

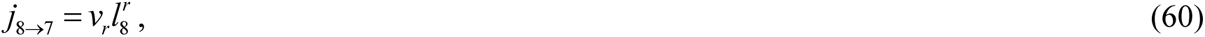

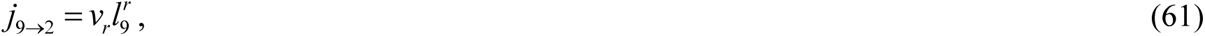

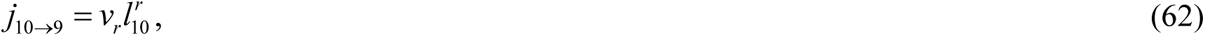

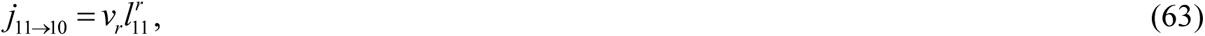

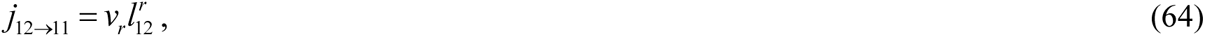

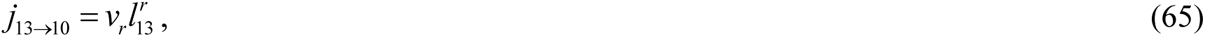

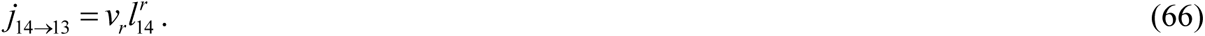

The turn-around fluxes (Fig. 2a) are simulated by the following equations:

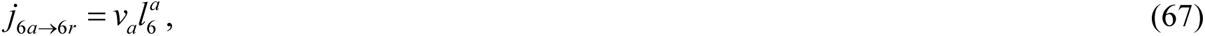

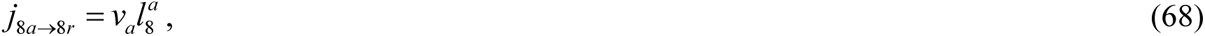

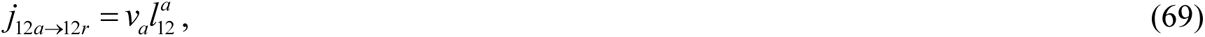

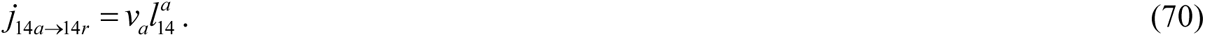

### 2.2. Model of mean age and age density distributions of mitochondria in the demand sites for the symmetric axon displayed in Figs. 1a and 2a

Although the governing equations (1)-(42) are easier to interpret in terms of the concentration of mitochondria in a compartment, to make further progress, they need to be re-scaled in the form given by Eqs. (71) and (72) below. This is because the conserved quantity is the total length of mitochondria in a compartment, not its linear density (concentration). Re-stating the governing equations in the form of Eqs. (71) and (72) enables the utilization of the methods developed in Metzler et al. (2018) and Metzler and Sierra (2018) for computing the system age, mean age, and age density distributions of mitochondria in various compartments.

Using the methodology described in Anderson (1983), Eqs. (1)-(42) were transformed into the following matrix equation:

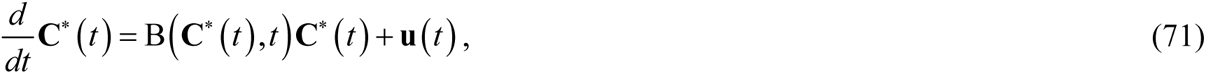

where **C**\* is referred to as a state vector (using the terminology of Metzler et al. 2018). **C**\* is defined by its components:

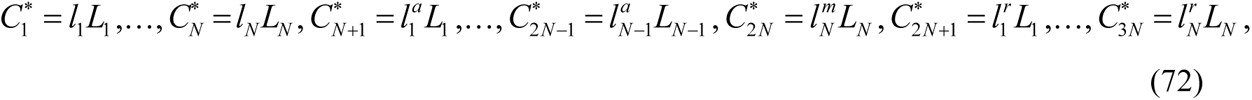

The total length of mitochondria in stationary states is represented by the first *N* components of the state vector **C**\*, the total length of mitochondria in anterograde states is represented by the next *N* components (*N*+1,…,2*N*), and the total length of mitochondria in retrograde states is represented by the last *N* components (2*N*+1,…,3*N*), see Fig. 2a.

In Eq. (72)

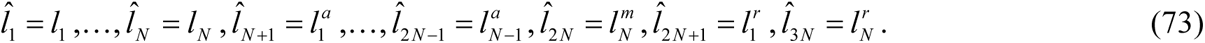

Vector **u**(*t*) in Eq. (71) is defined below by Eqs. (202) and (203).

We accounted for the internal mitochondrial fluxes between the compartments, the external mitochondrial flux entering the terminal from the axon, and the mitochondrial flux leaving the terminal to move back to the axon (Fig. 2a). Equations for the following elements of matrix B defined in Eq. (71) were obtained by analyzing mitochondrial fluxes to and from the compartments surrounding the most proximal demand site (utilizing the method of developing the compartmental model equations described in Anderson 1983).

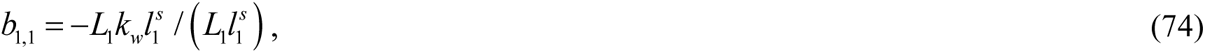

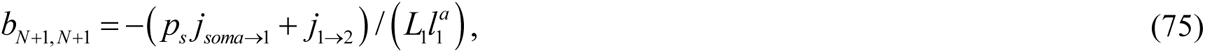

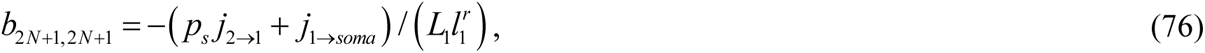

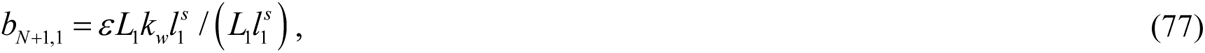

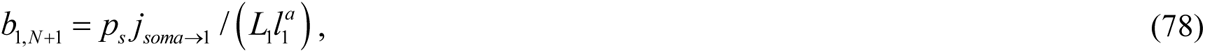

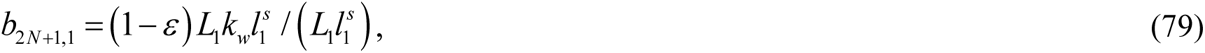

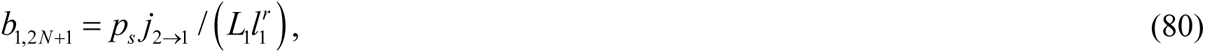

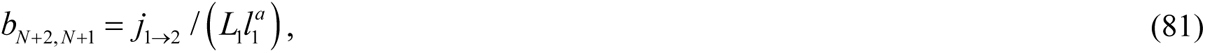

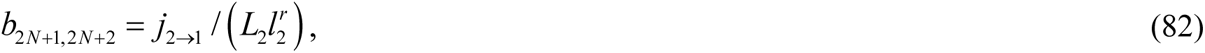

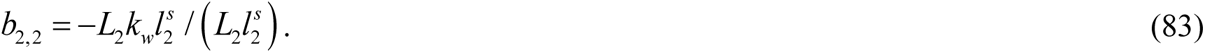

For other demand sites (again using the method described in Anderson 1983):

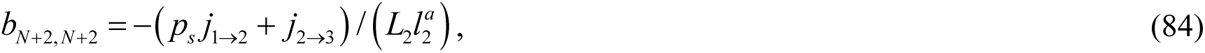

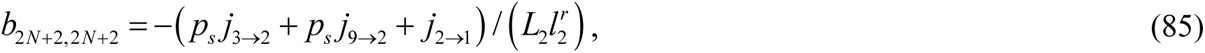

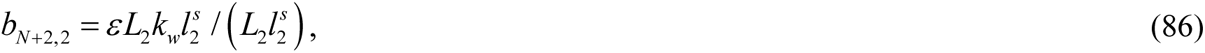

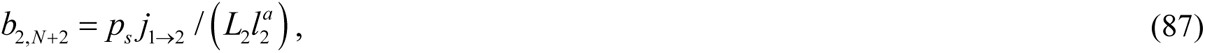

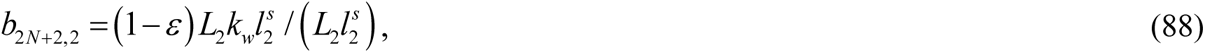

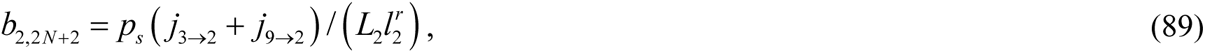

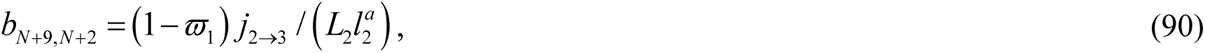

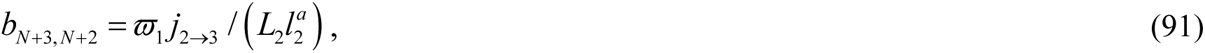

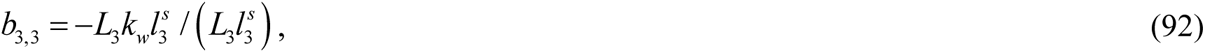

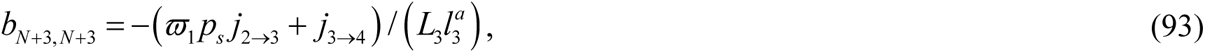

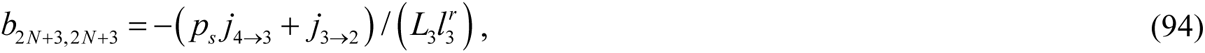

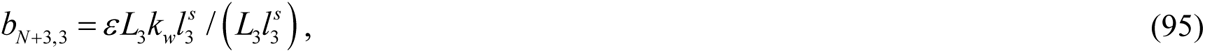

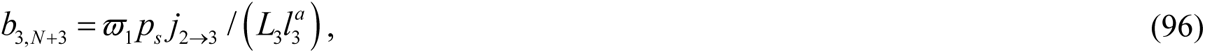

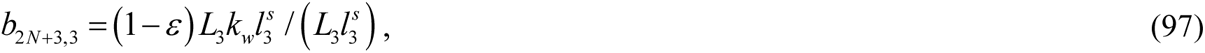

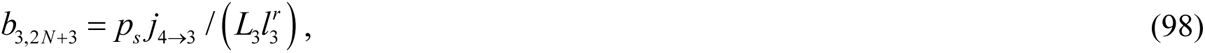

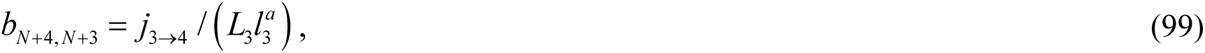

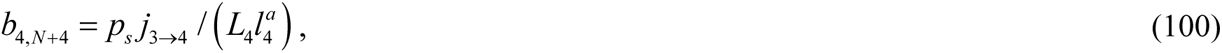

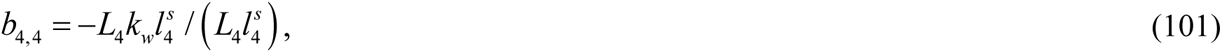

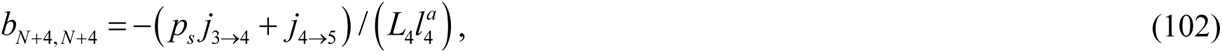

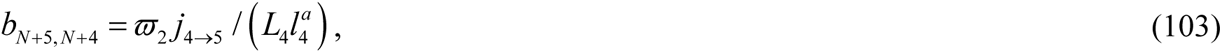

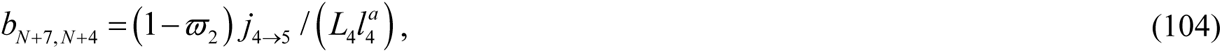

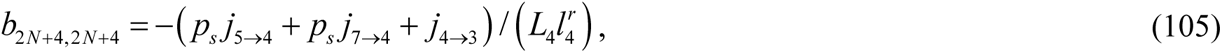

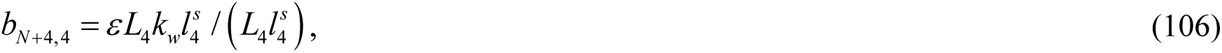

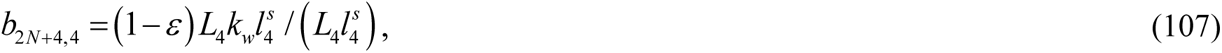

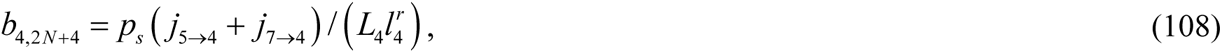

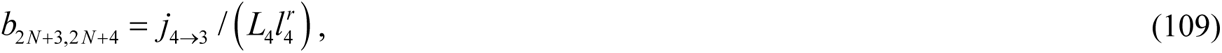

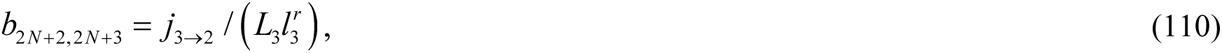

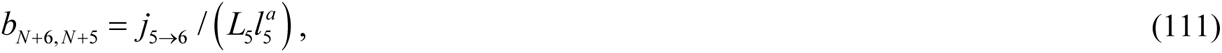

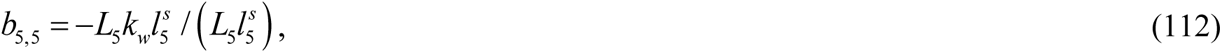

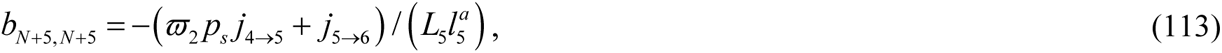

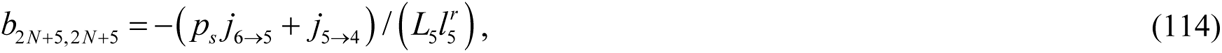

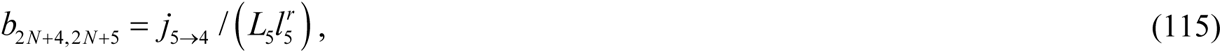

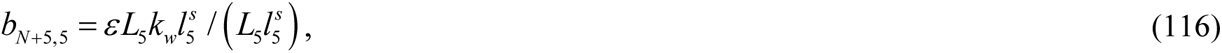

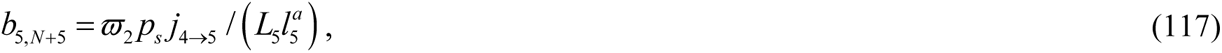

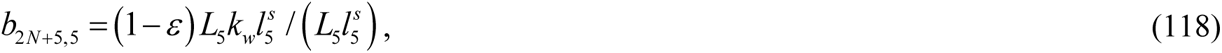

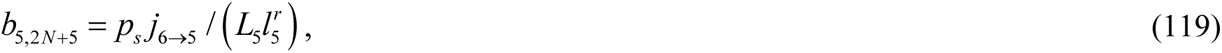

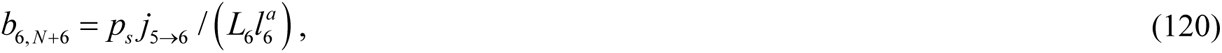

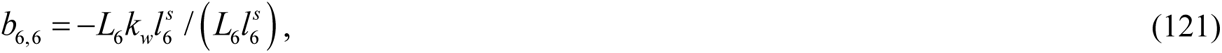

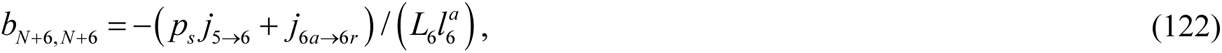

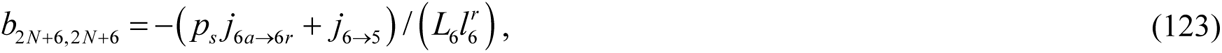

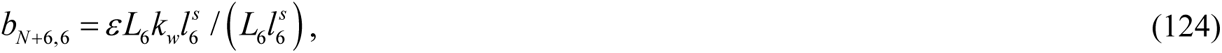

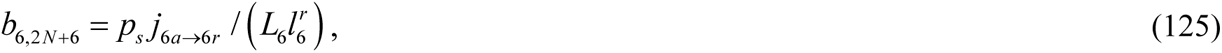

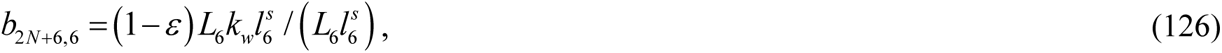

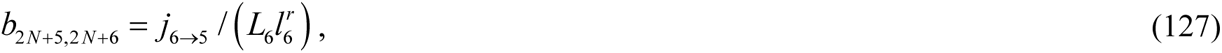

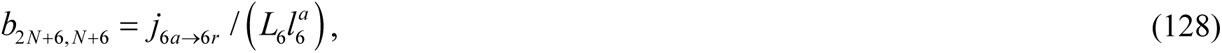

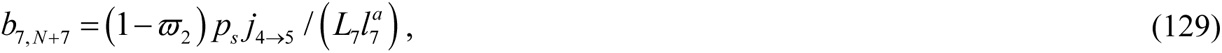

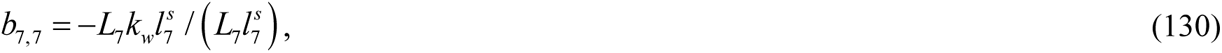

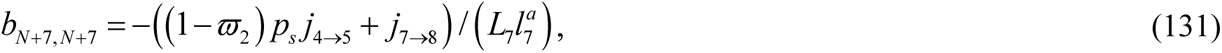

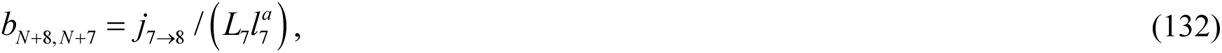

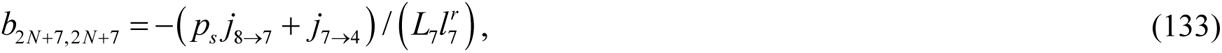

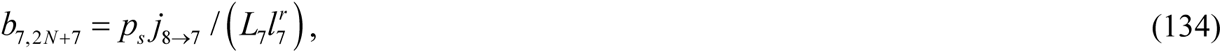

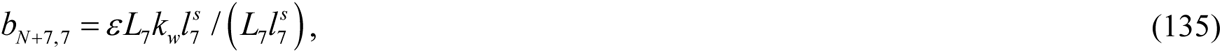

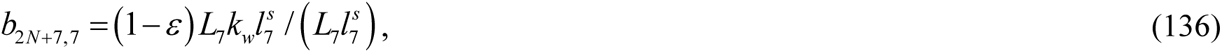

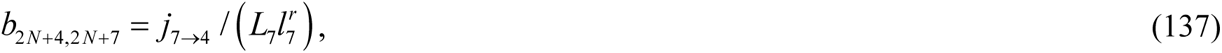

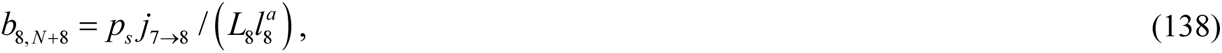

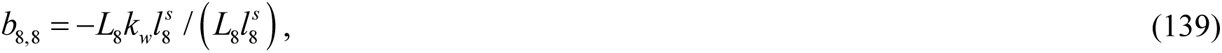

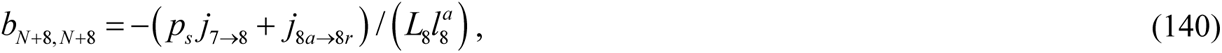

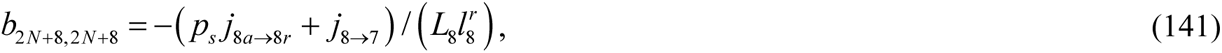

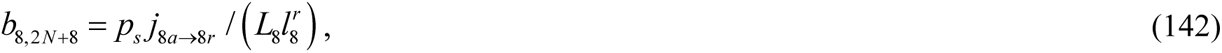

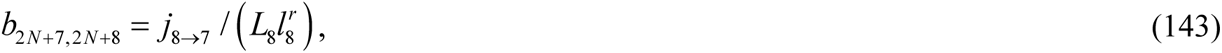

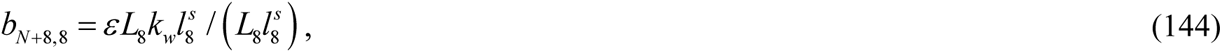

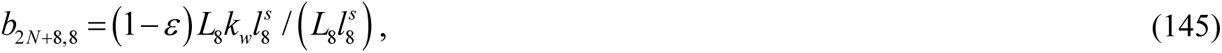

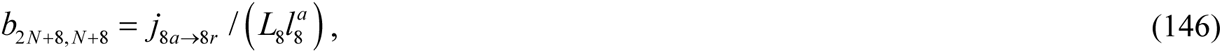

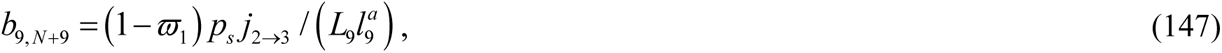

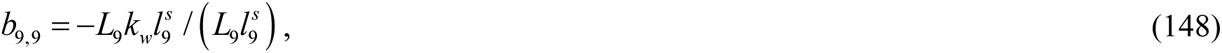

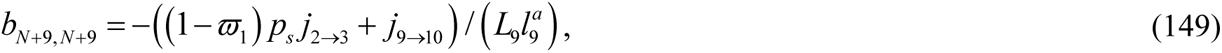

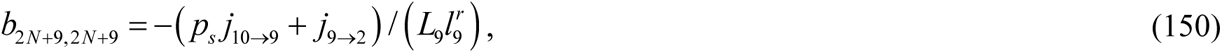

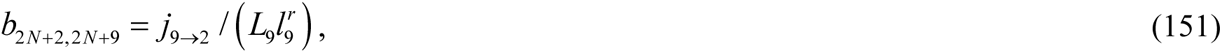

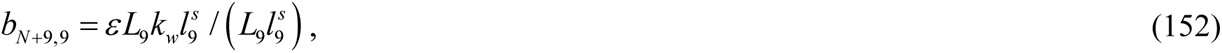

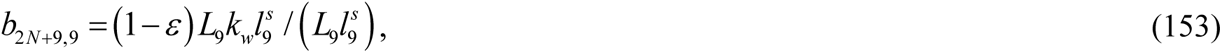

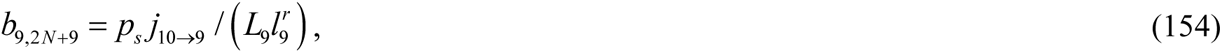

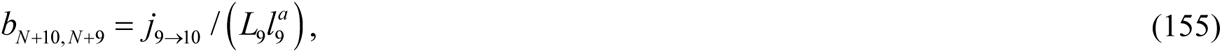

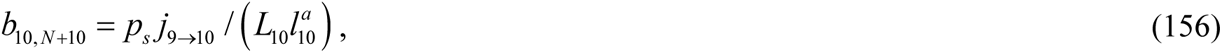

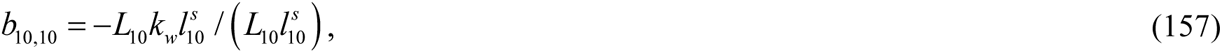

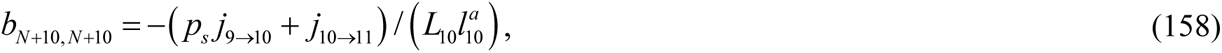

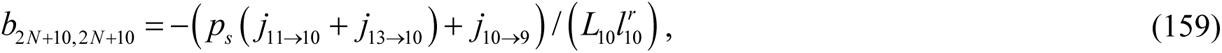

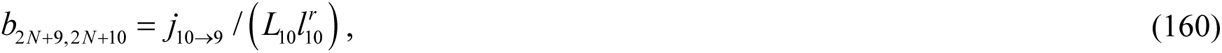

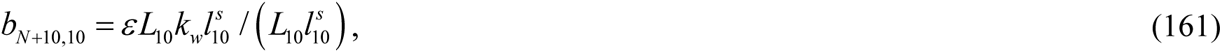

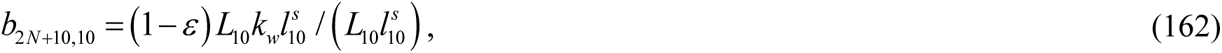

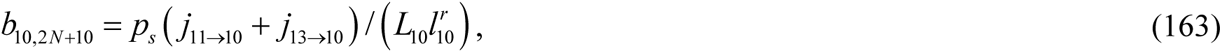

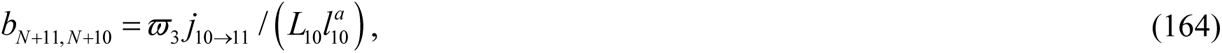

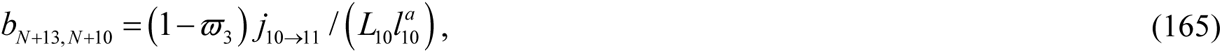

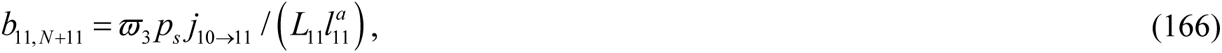

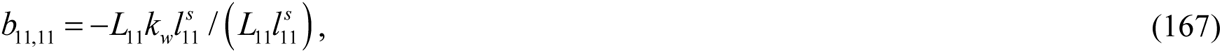

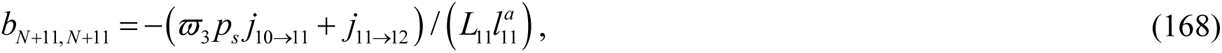

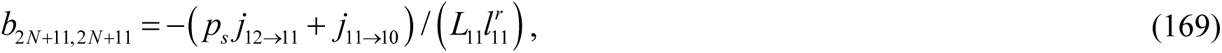

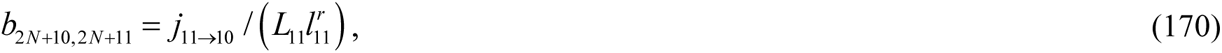

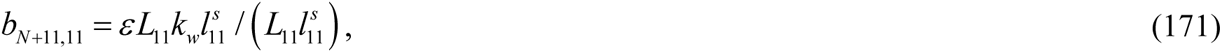

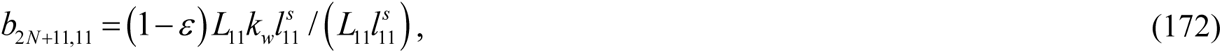

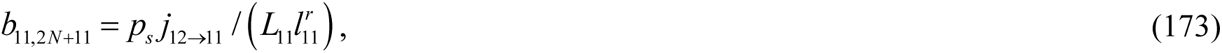

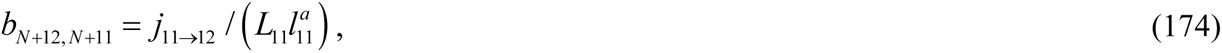

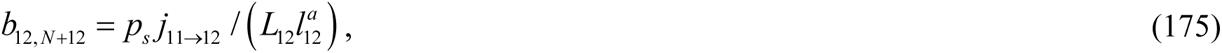

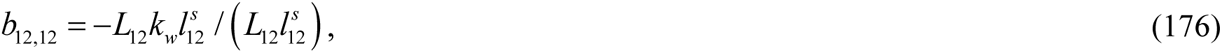

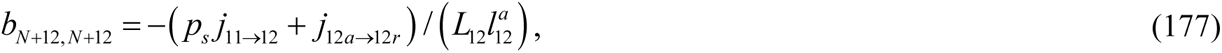

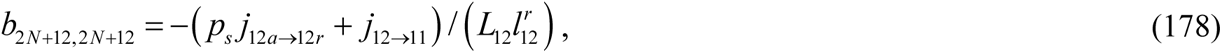

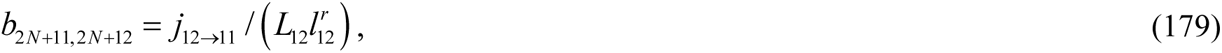

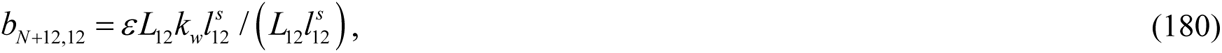

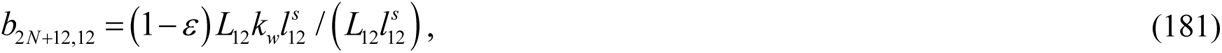

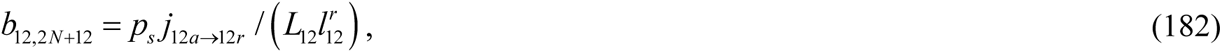

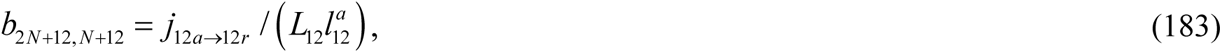

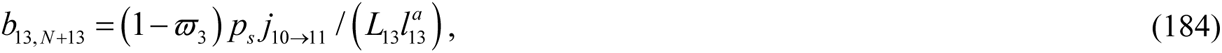

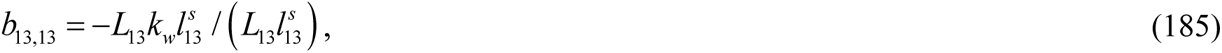

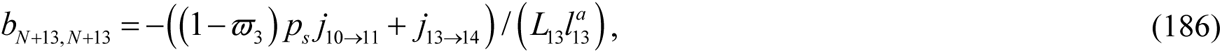

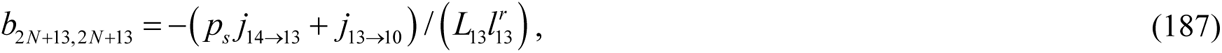

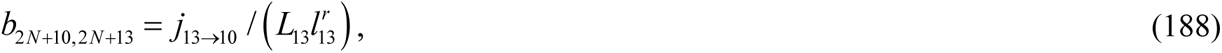

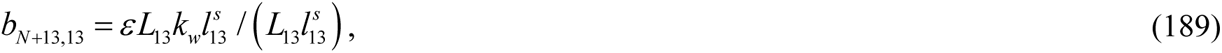

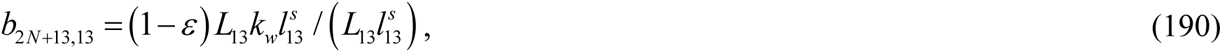

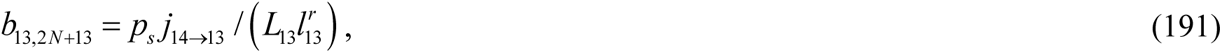

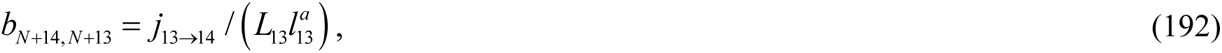

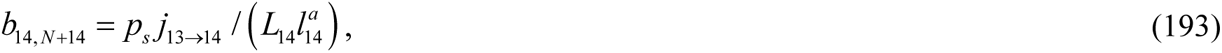

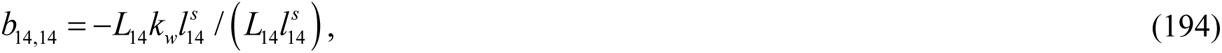

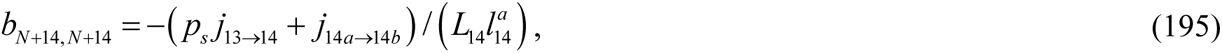

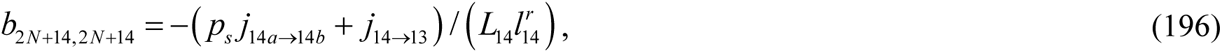

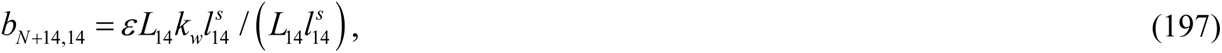

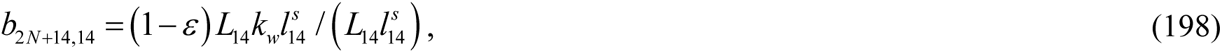

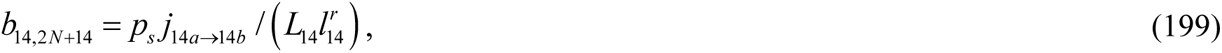

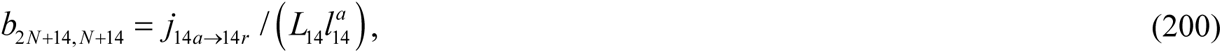

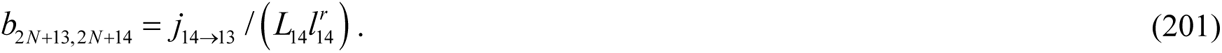

All other elements of matrix B, apart from those given by Eqs. (74)-(201), are equal to zero. When considering a single mitochondrion in the axon (its behavior is independent of all other mitochondria), the elements on the main diagonal determine the length of time it remains in a given compartment, while the off-diagonal elements describe the probabilities of it transitioning to other compartments (Metzler and Sierra 2018).

The only flux entering the axon is the anterograde flux from the soma to the compartment with anterograde mitochondria by the most proximal demand site, *j_soma_* _→1_. Our model assumes that any mitochondria leaving the axon (their flux is *j*_1→_*_soma_*) are degraded in the soma and not re-entering the axon. We also assume that all mitochondria entering the axon (their flux is *j_soma_* _→1_) are newly synthesized in the soma and have an age of zero at the time of entry. The mitochondrial age is calculated as the time since the mitochondria entered the axon.

The following equation yields the (*N*+1)^th^ element of vector **u**, which represents mitochondrial flux entering the axon from the soma:

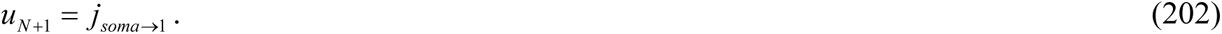

As no other external fluxes enter the terminal,

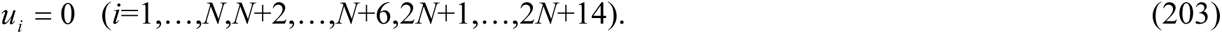

### 2.3. Calculating the mean age and age density distribution of mitochondria

A steady state solution for Eq. (71) can be obtained as follows (Table 1 of Metzler and Sierra 2018):

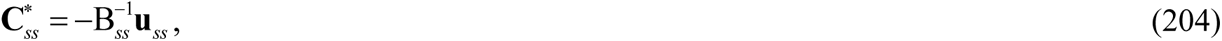

where the superscript −1 on a matrix denotes the inverse of the matrix, while the subscript *ss* stands for steady state. The mean ages of mitochondria in various compartments displayed in Fig. 2a at steady state can be calculated as follows:

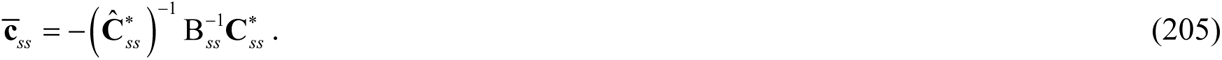

In Eq. (205)

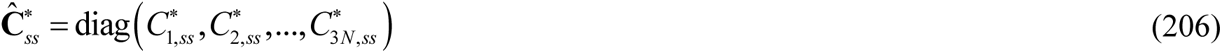

is the diagonal matrix with the components of the state vector, which is defined in Eq. (72), on the main diagonal. These components (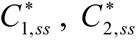, etc.) are determined at the steady state. Furthermore, in Eq. (205)

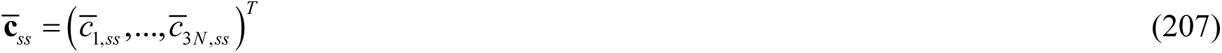

represents a column vector composed of the mean ages of mitochondria in various compartments (superscript *T* means transposed).

The vector whose components represent the age density of mitochondria residing in various compartments at steady state can be calculated as follows:

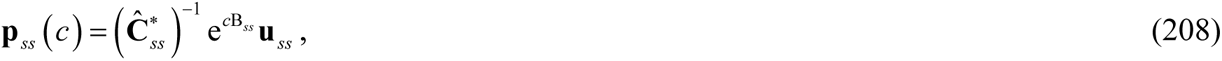

where e denotes the matrix exponential. The age density of mitochondria can be defined as the proportion of mitochondria that have an age within a specific interval, *T* and *T*+*dT*, throughout *dT*. The total number of mitochondria in a particular kinetic state can be obtained by integrating the age density of mitochondria from 0 to infinity. Additionally, integrating over any time interval will yield the number of mitochondria that fall within that time interval.

The system age is defined as the average age of all mitochondria across the entire axon and is not equivalent to the compartmental mean age defined by Eq. (205). The system age is calculated as follows (Metzler and Sierra 2018; Ceballos-Núñez et al. 2018):

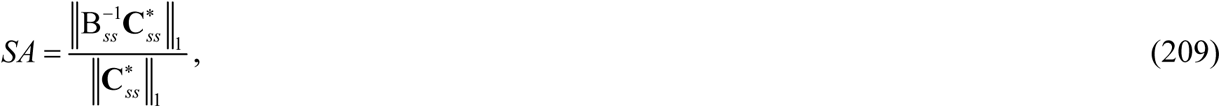

where the 1-norm of vector 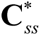 (for example) is defined as follows:

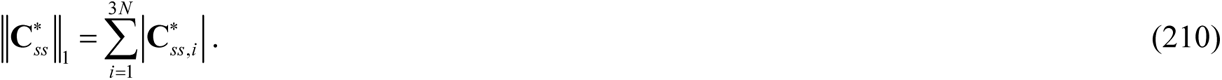

System age density can be calculated as follows (Metzler and Sierra 2018):

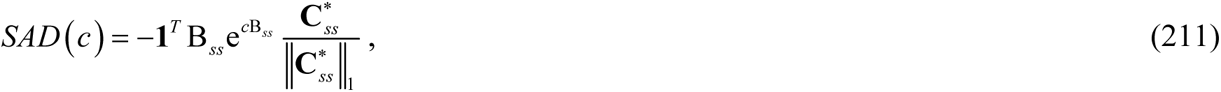

where **1***^T^* is the 3*N*-dimensional row vector whose entries are equal to one. Because *SAD* is a probability density function,

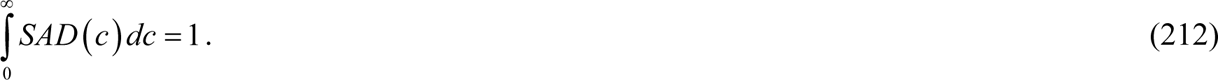

Governing equations for the case of an axon with two asymmetric branches (Figs. 1b and 2b) are given in section S1 in Supplemental Materials.

### 2.4. Numerical solution procedure

In this paper, we utilized two different methods for computing the steady state distributions of mitochondria in the axon. These two methods were used to validate each other. The first method involves solving the transient equations until the numerical solutions reach a steady state. Since we are only interested in the solution at steady state, we can choose any initial condition (for simplicity, we used the zero boundary condition). Eqs. (1)-(42), which simulate the transport of mitochondria in the axon with two branches of equal length (as shown in Fig. 1a), were solved using the MATLAB solver ODE45 (MATLAB R2020b, MathWorks, Natick, MA, USA).

Eqs. (204)-(208) allow us to find steady state solutions directly. We implemented these equations using standard MATLAB operators, such as matrix inverse and matrix exponential.

## 3. Results

The calculations reported in Figs. 3-7 and S1-S6 in this paper are performed for the axons depicted in Fig. 1a and 1b, considering the scenario where the flow of mitochondria is divided equally at each branching junction (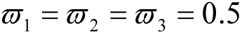 for the axon depicted in Fig. 1a and 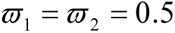 for the axon depicted in Fig. 1b). As a result, the concentration of mitochondria decreases by half at each branching junction (Figs. 3a and 3b). This is due to the decrease in the rate of flow of mitochondria, which is proportional to the concentration of mitochondria (as described in Eqs. (43)-(70) and (S31)-(S50)). Note that the effect of mitophagy was neglected. The mean age of anterogradely moving mitochondria in the symmetric (Fig. 1a) and asymmetric (Fig. 1b) axons increases from the most proximal (#1) to the most distal (#6) demand site, i.e. 1 → 2 → 3 → 4 → 5 → 6 (Fig. 4). This is because the mitochondria get older as they move from one demand site to the next (see the age density distributions displayed in Figs. S1-S6). The same holds true for stationary mitochondria. The pattern for retrograde mitochondria is more complicated, as two opposing trends shape their distribution: (i) they get older as they move from the more proximal to the more distal demand site, and (ii) there is a transfer of newer mitochondria from stationary states (Fig. 2a,b). Fig. 4 shows that in the asymmetric axon (Fig. 1b), the mean ages of mitochondria are lower due to its shorter total branch length compared to the symmetric axon (Fig. 1a). The average ages of mitochondria in the axon (system age) calculated according to Eq. (209) for the axon with symmetric branches (Fig. 1a) and for the axon with asymmetric branches (Fig. 1b) are given in Table 3.

**Fig. 3.**
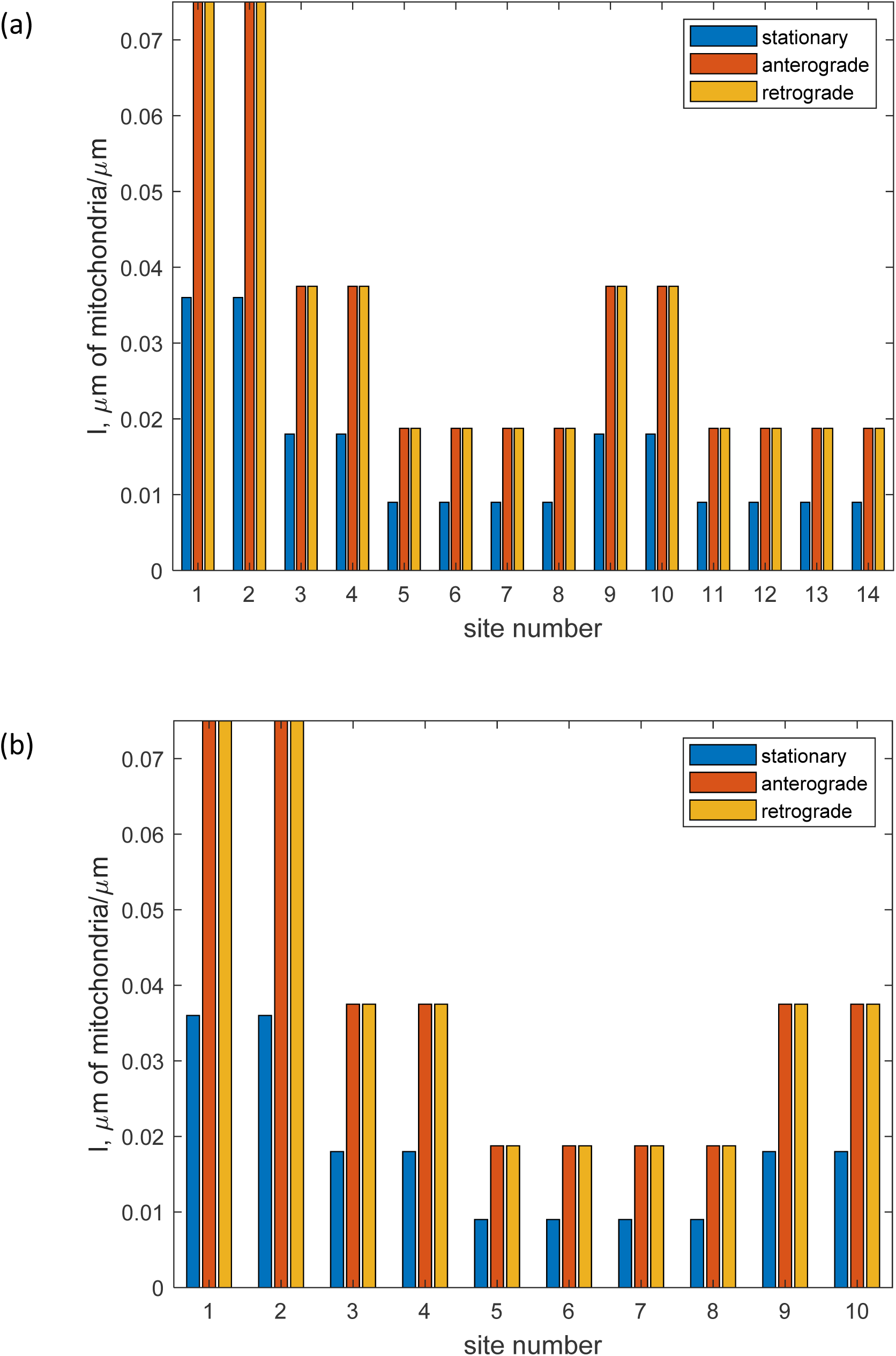
The steady state values of the total length of stationary, anterogradely moving, and retrogradely moving mitochondria per unit length of the axon in the *i*-th demand site’s compartment. (a) shows the results for a symmetric branched axon, and (b) displays the results for an asymmetric branched axon. *ω*_1_ = *ω*_2_ = *ω*_3_ = 0.5. In this case, the mitochondrial flux splits in half at each branching junction, and the concentration of mitochondria reduces by 50%.

**Table 3.** System age of mitochondria in the axon (calculated according to Eq. (209)).

|  | Axon with symmetric branches (Fig. 1a) | Axon with asymmetric branches (Fig. 1b) |
| --- | --- | --- |
| $\omega_1 = \omega_2 = \omega_3 = 0.5$ (symmetric axon),<br>$\omega_1 = \omega_2 = 0.5$ (asymmetric axon) | 12.13 hours | 10.04 hours |
| $\omega_1 = \omega_2 = \omega_3 = 0.9$ (symmetric axon),<br>$\omega_1 = \omega_2 = 0.9$ (asymmetric axon) | 12.13 hours | 11.75 hours |

The ages of mitochondria given in Table 3 and plotted in Fig. 4 are comparable with the estimate reported in Kuznetsov and Kuznetsov (2022a) (22 hours). Kuznetsov and Kuznetsov (2022a) estimated the mean age of mitochondria using experimental data reported in Ferree et al. (2013). The results in Table 3 and Fig. 4 are also consistent with Mandal et al. (2021), which reported that the complete turnover of axon terminal mitochondria in zebrafish neurons occurs within 24 hours.

**Fig. 4.**
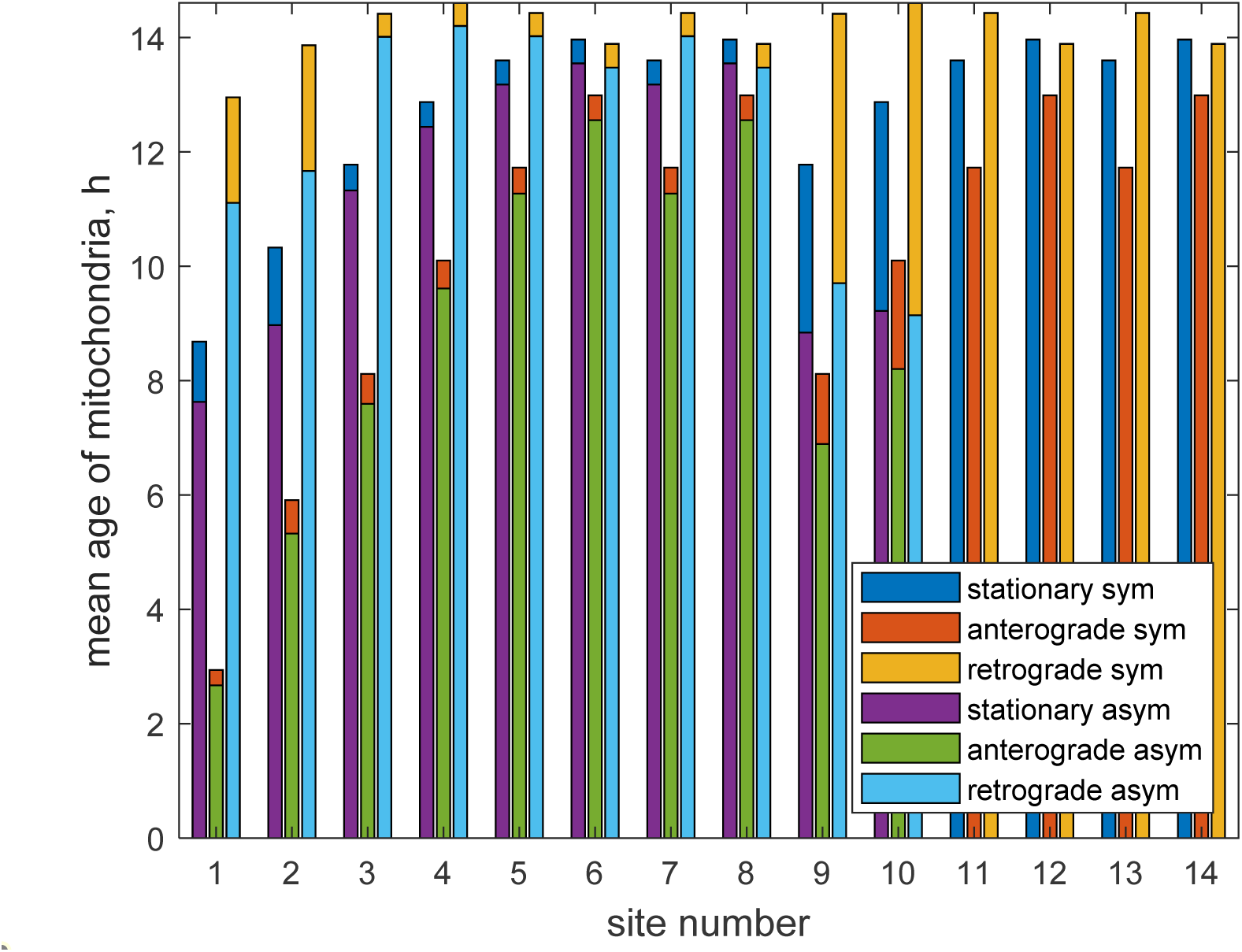
Comparison of the mean age of stationary, anterogradely moving, and retrogradely moving mitochondria in symmetric (a) and asymmetric (b) branched axons. *ω*_1_ = *ω*_2_ = *ω*_3_ = 0.5. Due to its shorter total branch length, the mean ages of mitochondria in the asymmetric axon are lower than in the symmetric axon.

The mean ages of mitochondria do not provide a comprehensive understanding of the age of mitochondria within the axon. Each demand site contains mitochondria of different ages, as illustrated by the mitochondrial age density distributions in symmetric (Figs. S1-S3) and asymmetric (Figs. S4-S6) axons. The presence of long tails suggests that each demand site contains mitochondria that are significantly older than the mean age.

The newly formed anterograde mitochondria with a zero-age are delivered to the anterograde compartment at the most proximal demand site (#1). This is why the highest concentration of anterograde mitochondria is found at zero-age at the most proximal demand site (Fig. 5a). As the demand sites get farther away from the soma, the peak of the age density distribution for anterograde mitochondria shifts towards older mitochondria (Fig. 5a). The age density distributions of mitochondria in demand sites 8 and 14 in the axon shown in Fig. 1a are the same due to symmetry (Figs. 5b, 6b, and 7b). However, the age density distribution at demand site 10 is smaller due to its proximity to the soma compared to demand site 8 (Fig. 1a, 1b).

**Fig. 5.**
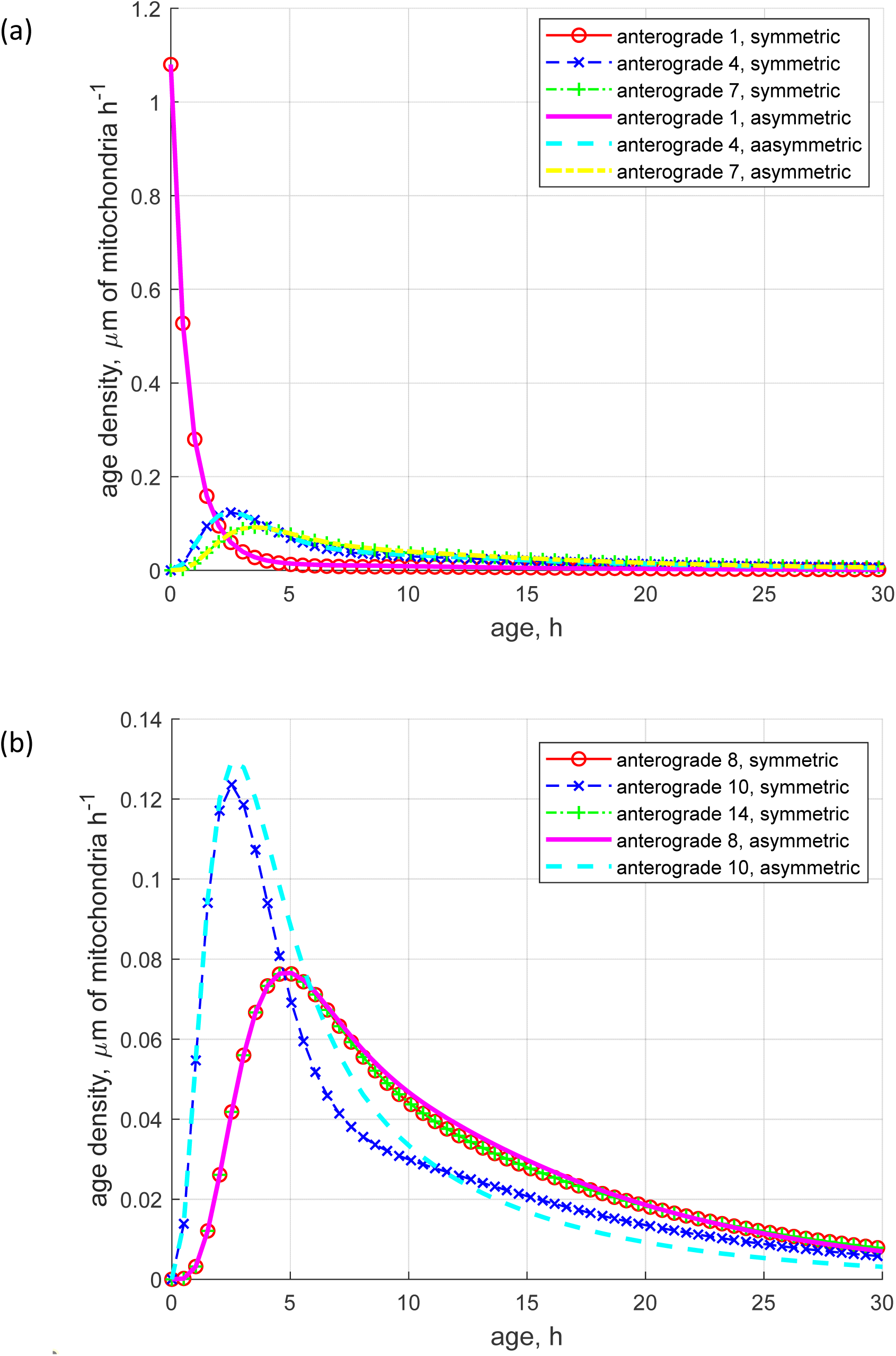
Comparison of the age density distributions of anterogradely moving mitochondria in symmetric and asymmetric branched axons in various demand sites. (a) Demand sites 1, 4, 7. (b) Demand sites 8, 10, 14 (14 only for the symmetric axon). *ω*_1_ = *ω*_2_ = *ω*_3_ = 0.5. The highest concentration of newly formed anterograde mitochondria occurs at zero-age at the most proximal demand site, and the peak of the age density distribution for anterograde mitochondria shifts towards older mitochondria as the demand sites get farther away from the soma.

**Fig. 6.**
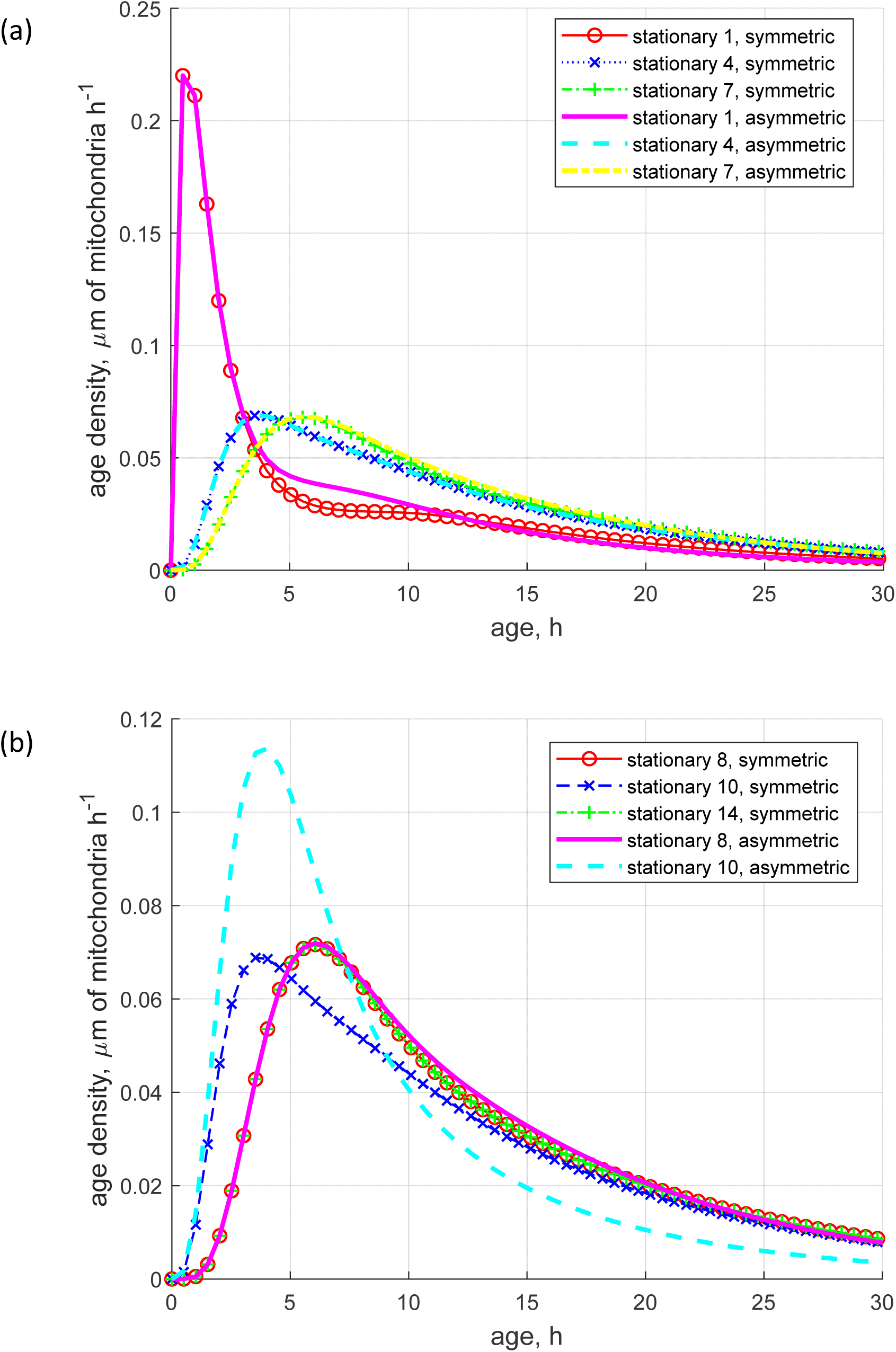
Comparison of the age density distributions of stationary mitochondria in symmetric and asymmetric branched axons in various demand sites. (a) Demand sites 1, 4, 7. (b) Demand sites 8, 10, 14 (14 only for the symmetric axon). *ω*_1_ = *ω*_2_ = *ω*_3_ = 0.5. The discrepancy between the age density distributions in symmetric and asymmetric axons is more noticeable for stationary mitochondria than for anterograde mitochondria.

The difference between the age density of anterograde mitochondria in symmetric and asymmetric axons is not very visible (Fig. 5). The difference in mean ages seen in Fig. 4 primarily results from the skewed right shape of the age density distributions and longer tails in the positive direction for the symmetric axon. This indicates a higher number of older mitochondria in the symmetric axon (Metzler and Sierra 2018). This is because the symmetric axon (Fig. 1a) has longer branches than the asymmetric axon (Fig. 1b), which leads to a longer travel time for retrograde mitochondria. However, the difference between the age density distributions in symmetric and asymmetric axons becomes more pronounced for stationary mitochondria (compare the age density distributions for demand site 1 in Fig. 6a) and particularly for retrograde mitochondria (compare the age density distributions for demand site 1 in Fig. 7a).

**Fig. 7.**
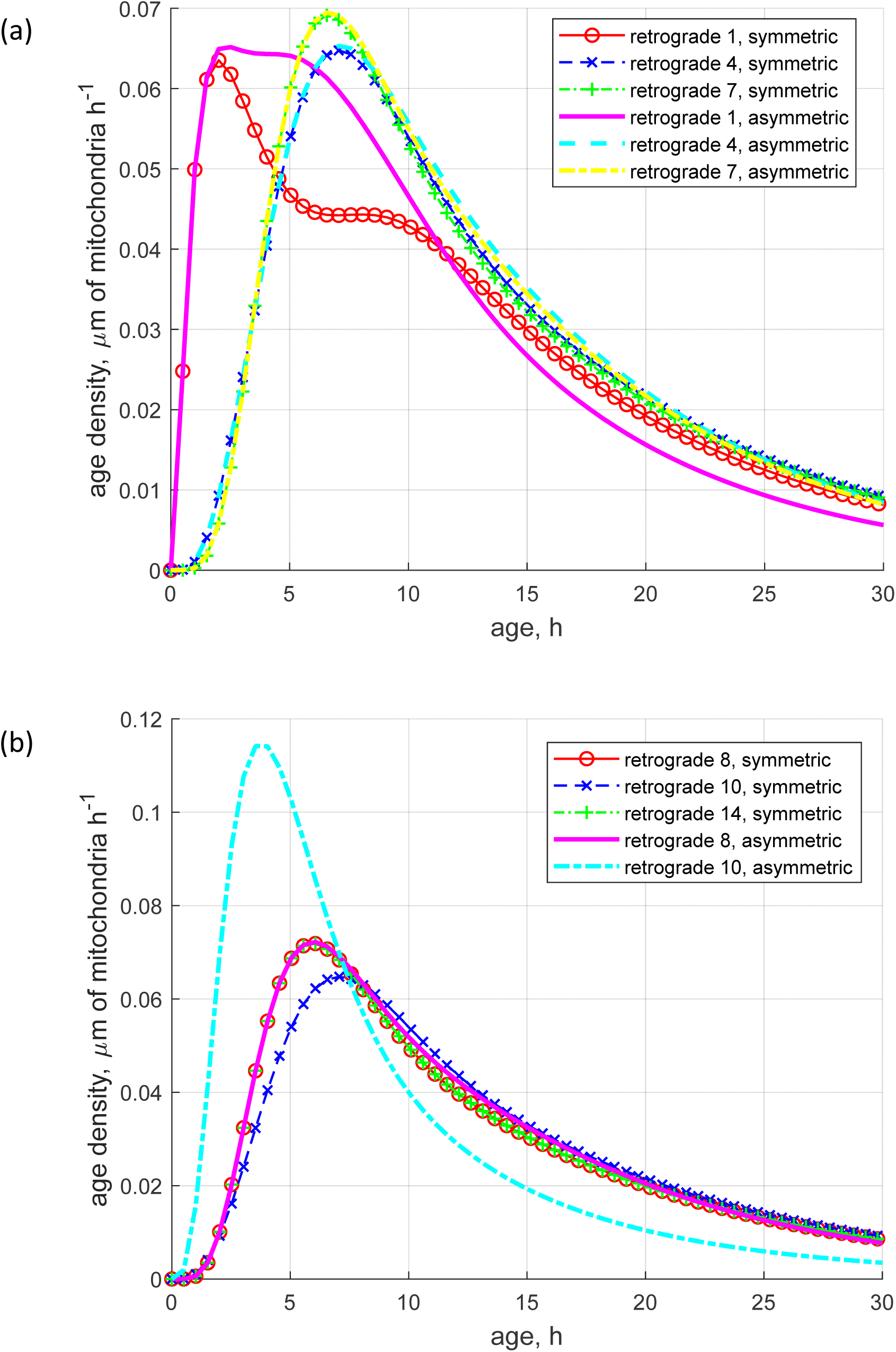
Comparison of the age density distributions of retrograde moving mitochondria in symmetric and asymmetric branched axons in various demand sites. (a) Demand sites 1, 4, 7. (b) Demand sites 8, 10, 14 (14 only for the symmetric axon). *ω*_1_ = *ω*_2_ = *ω*_3_ = 0.5. The difference between the age density distributions in symmetric and asymmetric axons is even greater for retrograde mitochondria.

Recent research indicates that the anterograde transport of organelles through branching junctions is selective and tightly regulated (Tymanskyj et al. 2022). In our model, the way the mitochondrial flux splits at the branching junctions is controlled by parameters *ω*_1_, *ω*_2_, and *ω*_3_ (Fig. 1a,b). The increase in the values of *ω*_1_, *ω*_2_, and *ω*_3_ to 0.9 results in a much larger mitochondrial flux entering the upper branches at the branching junction. As a result, mitochondria concentrations in the upper branches increase proportionally (Fig. 8). It should be noted that a greater mitochondrial flux into a longer branch is expected because a longer branch has more demand sites and hence needs more mitochondria.

**Fig. 8.**
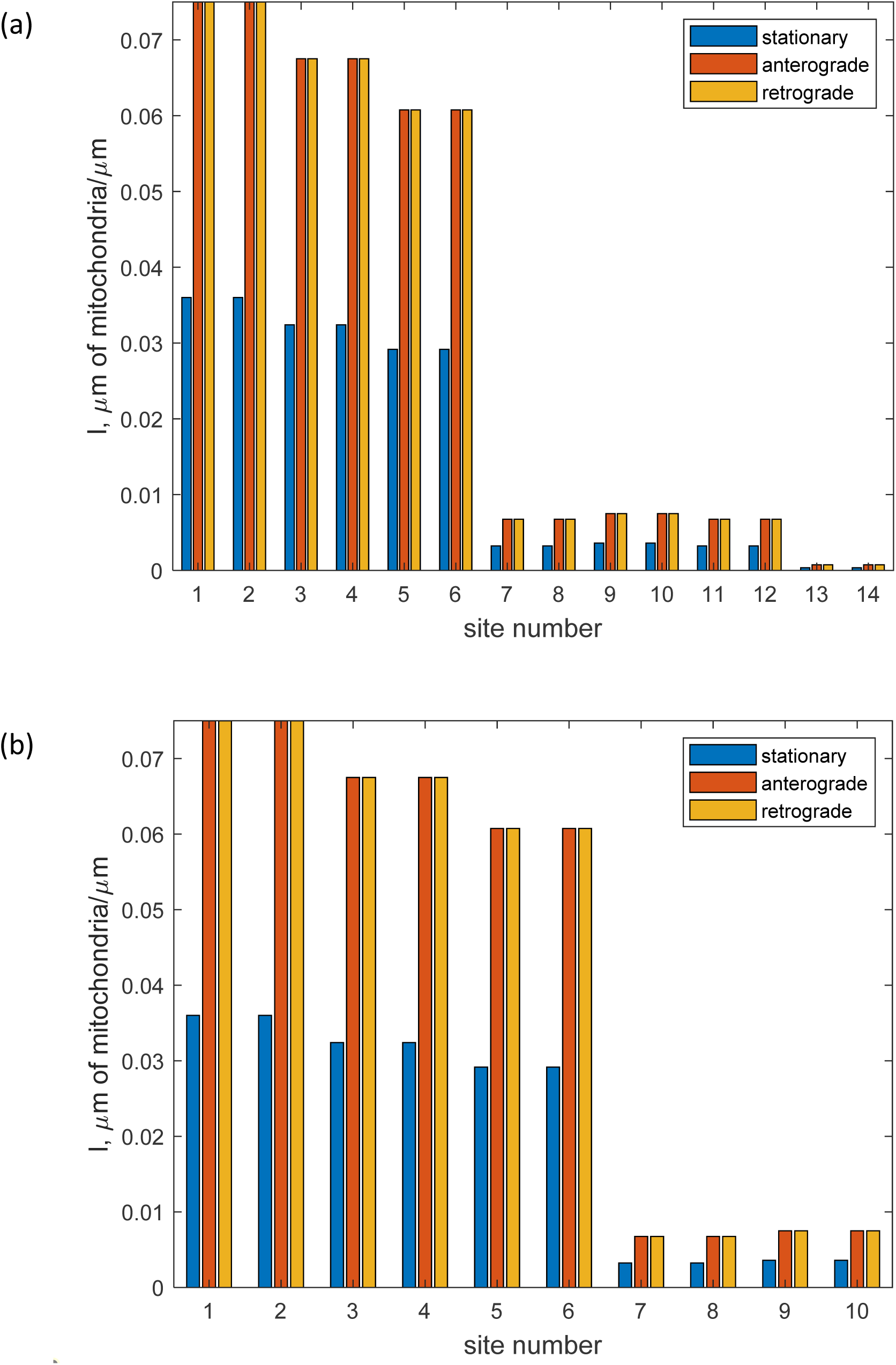
The steady state values of the total length of stationary, anterogradely moving, and retrogradely moving mitochondria per unit length of the axon in the *i*-th demand site’s compartment. (a) shows the results for a symmetric branched axon, and (b) displays the results for an asymmetric branched axon. *ω*_1_ = *ω*_2_ = *ω*_3_ = 0.9. Increasing the values of *ω* -s to 0.9 leads to a significant increase in the flux of mitochondria entering the upper branches at the branching junction. This, in turn, causes an increase in the concentrations of mitochondria in the upper branches.

The system age density for the symmetric axon is independent of how the flux of mitochondria splits at the branching junction (Fig. 9a). Consequently, the age densities of mitochondria in anterograde, stationary, and retrograde compartments by various demand sites in the symmetric axon are independent of the values of *ω* -s (results not shown). This observation can be easily explained using the one-particle perspective (Metzler and Sierra 2018). In our model, the assumption is made that the movement of mitochondria along the axon is not affected by the presence of other mitochondria. Consequently, the age of a particular mitochondrion is not determined by the percentage of mitochondria that have taken a specific branch at the branching junction as long as the branches are of the same length.

**Fig. 9.**
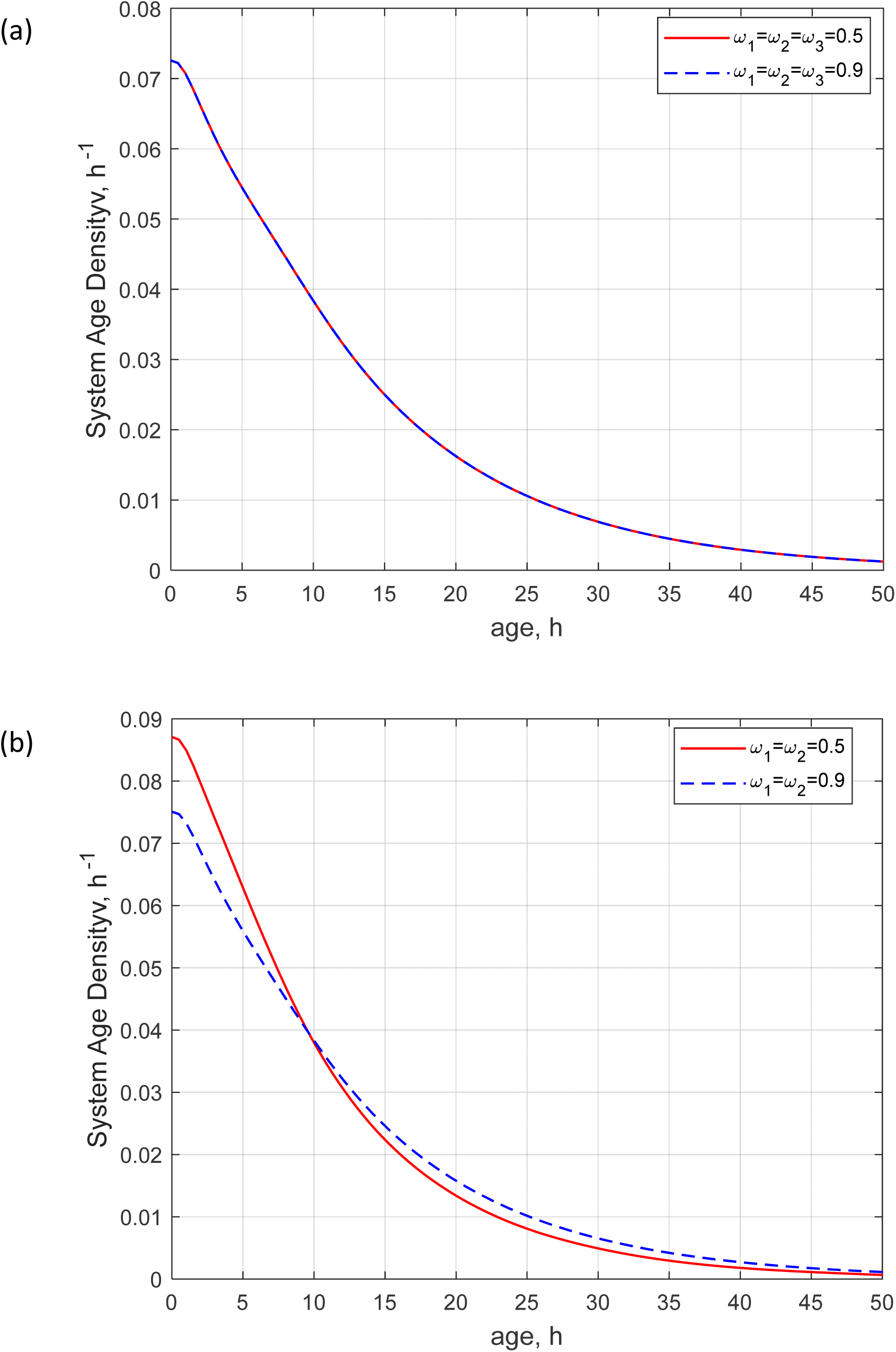
System age density. (a) Comparison of the cases *ω*_1_ = *ω*_2_ = *ω*_3_ = 0.5 and *ω*_1_ = *ω*_2_ = *ω*_3_ = 0.9 for the symmetric axon displayed in Fig. 1a. (b) Comparison of the cases *ω*_1_ = *ω*_2_ = 0.5 and *ω*_1_ = *ω*_2_ = 0.9 for the asymmetric axon displayed in Fig. 1b. The distribution of system age density in the symmetric axon remains constant regardless of how the mitochondrial flux splits at the branching junction. In contrast, in the asymmetric axon, if more mitochondria enter the longer branch, the system age density of older mitochondria becomes higher.

However, for the asymmetric axon, the system age density of older mitochondria is larger when more mitochondria enter the longer branch (this corresponds to the case of *ω*_1_ = *ω*_2_ = 0.9 in Fig. 9b). This case is investigated in Figs. S7-S9, which compare age densities of mitochondria in anterograde (Fig. S7), stationary (Fig. S8), and retrograde (Fig. S9) compartments by various demand sites in the asymmetric axon. There is a notable difference between the age densities of anterogradely moving mitochondria between *ω*_1_ = *ω*_2_ = 0.5 and *ω*_1_ = *ω*_2_ = 0.9 cases in demand sites 9 and 10 (Fig. S7), the age densities of stationary mitochondria in demand sites 1, 2, and 3 (Fig. S8), and the age densities of retrogradely moving mitochondria in demand sites 1, 2, and 9 (Fig. S9). In all these demand sites, the age densities of older mitochondria are greater for the case of *ω*_1_ = *ω*_2_ = 0.9 than for the case of *ω*_1_ = *ω*_2_ = 0.5 because for the case of *ω*_1_ = *ω*_2_ = 0.9 more mitochondria enter the upper (longer) branch (Fig. 1b). This is consistent with the system age of mitochondria reported in Table 3.

## 4. Discussion, limitations of the model, and future directions

The average total length of dopaminergic neurons in rats is around 0.5 meters (Matsuda et al. 2009), which roughly corresponds to a neuron with 14 branching levels (Pissadaki and Bolam 2013). Our findings in this paper demonstrate the mitochondrial concentration decreases by half when the mitochondrial flux splits in half at each branching junction. This aligns with previous research indicating that the density of mitochondria is evenly divided at each branching junction (Agrawal and Koslover 2021). As a result, the mitochondrial concentration in the most distal branch is reduced by 2^14−1^ = 8192 times compared to the most proximal branch. The ATP concentration in axons is estimated to be between 2-4 mM (Pathak et al. 2015), and a minimum ATP concentration of 0.8 mM is needed in boutons to sustain synaptic transmission (Pathak et al. 2015). However, based on our findings, the ATP concentration in the most distal branch would be much lower at (2 − 4) mM / 8192 = (2.44 − 4.88) ×10^−4^ mM, which is far below the minimum required (0.8 mM). Further investigation is necessary to understand how dopaminergic neurons can maintain ATP concentration to sustain transmission in distal boutons.

Many neurons have branches of unequal length (Pissadaki and Bolam 2013; Duncan et al. 2018; Goaillard et al. 2020). We found that if the flux of mitochondria is split unevenly at the branching junction and a larger flux enters a longer branch, the system age density shifts toward older mitochondria. Since having branches with unequal length is typical for branching axons (Tymanskyj et al. 2022), this suggests a potential problem with maintaining uniform mitochondrial age (and, consequently, uniform mitochondrial health) that dopaminergic neurons with unequal branches may face. This observation may have relevance to PD since there is a potential association between mitochondrial dysfunction and the development of PD (Subramaniam and Chesselet 2013).

The developed model can simulate the time that mitochondria spend in compartments representing demand sites. This approach is valid for relatively short axons, such as one used for computations in this paper (we used an axon with a length of 1 cm, see Table 2). In the future, it is necessary to develop models that can estimate mitochondria age based on approaches that will account for the time it takes mitochondria to travel between demand sites, which can be significant for long axons. Another way to improve the current model is by preventing a scenario where newly entered mitochondria in the axon move quickly to the stationary state, followed by the retrograde state, and then exit the axon. This scenario, called short-circuiting (Bolin and Rodhe 1973), is unlikely to occur in real axons. To prevent this in the model, we can set the kinetic constants that describe the likelihood of mitochondria transitioning from the anterograde to the stationary state and then to the retrograde state for proximal demand sites to small values.

The current model neglects interactions between mitochondria. Crowding of mitochondria can cause congestion and even the formation of mitochondrial traffic jams (Correia et al. 2016). Extending the present model to add the capability to simulate the possible formation of mitochondrial traffic jams (as it was done for other organelles, see Kuznetsov and Avramenko (2009a, 2009b)) would be an interesting opportunity for future research.

## Acknowledgment

IAK acknowledges the fellowship support of the Paul and Daisy Soros Fellowship for New Americans and the NIH/National Institute of Mental Health (NIMH) Ruth L. Kirchstein NRSA (F30 MH122076-01). AVK acknowledges the support of the National Science Foundation (award CBET-2042834) and the Alexander von Humboldt Foundation through the Humboldt Research Award.

## Supplemental Materials

### S1. Governing equations for the case of an axon with two asymmetric branches (Figs. 1b and 2b)

#### S1.1. Equations that express the conservation of the total length of mitochondria in various compartments for an axon with two asymmetric branches (**Figs. 1b and 2b**)

Equations that are analogous to Eqs. (1)-(42) but now for the asymmetric axon displayed in Fig. 1b and 2b are

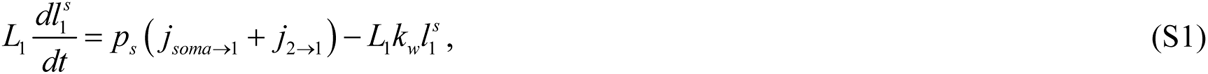

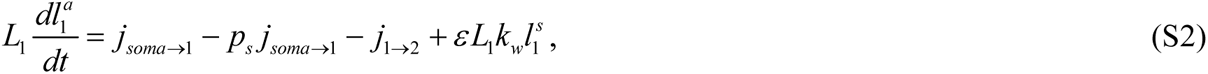

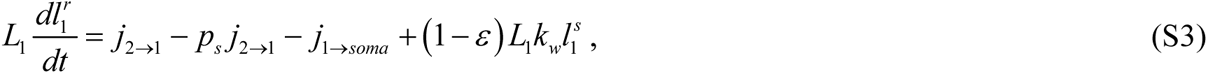

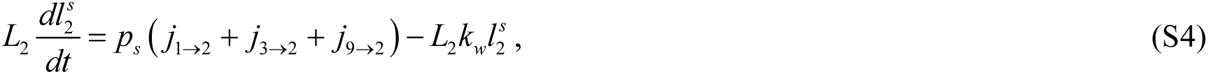

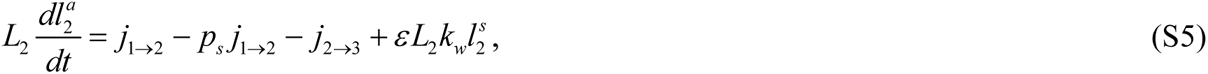

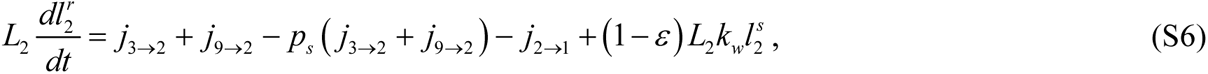

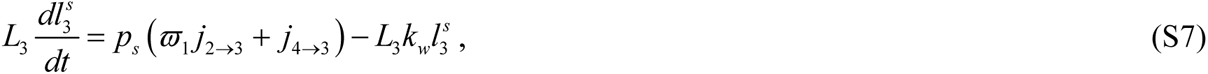

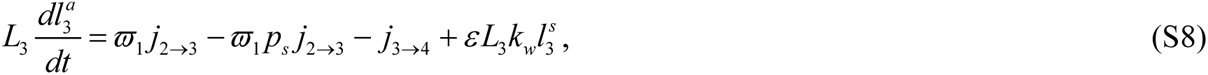

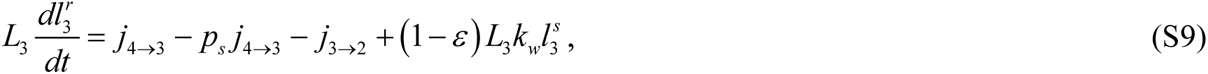

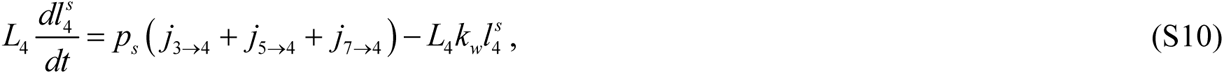

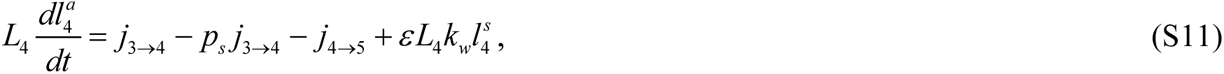

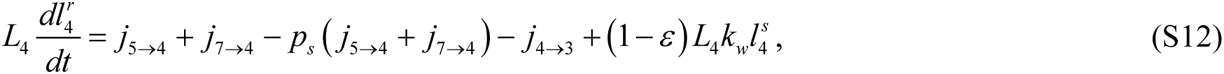

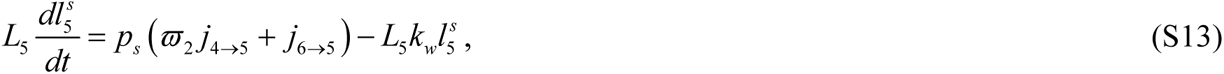

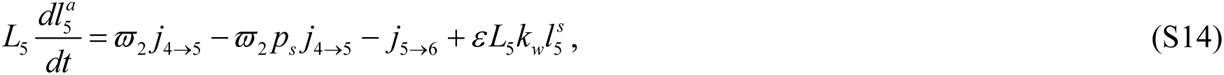

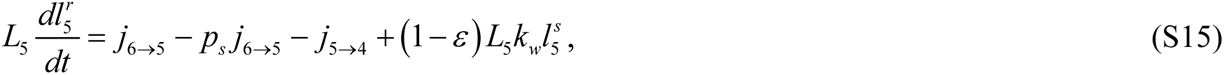

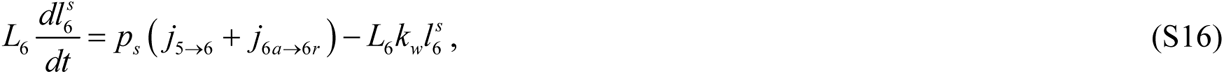

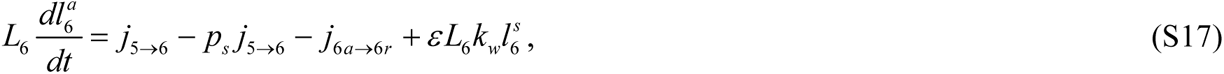

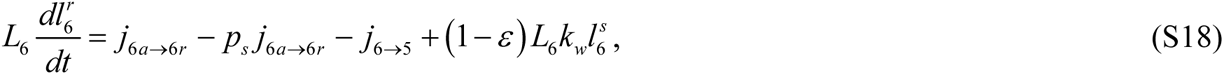

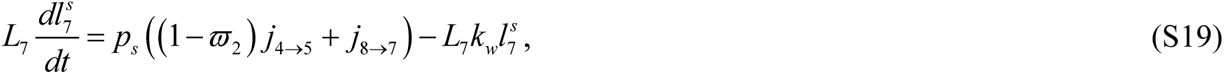

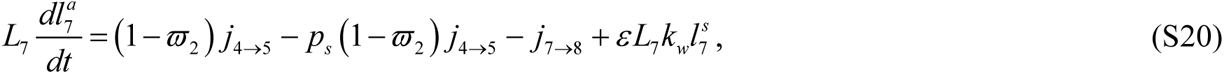

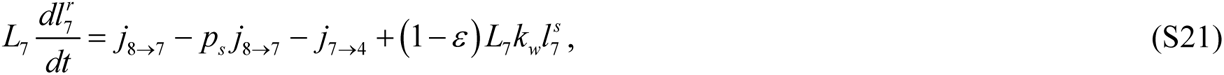

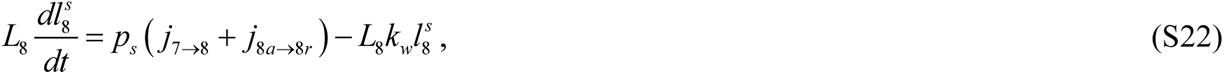

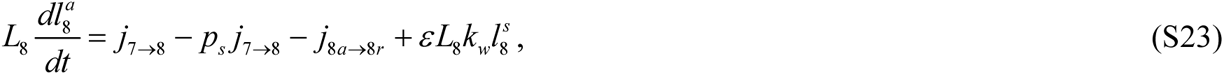

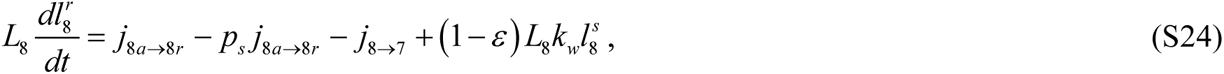

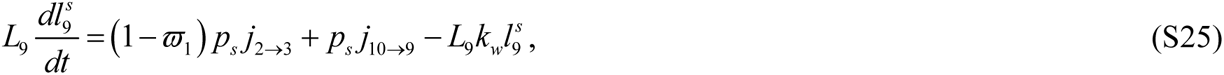

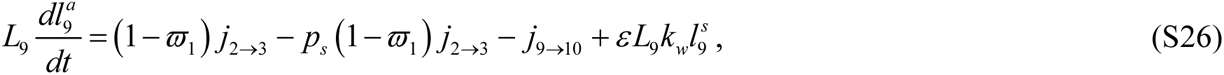

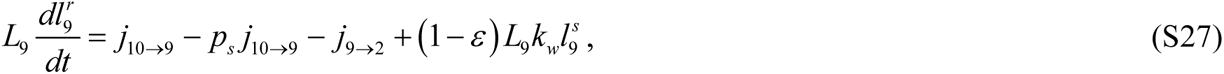

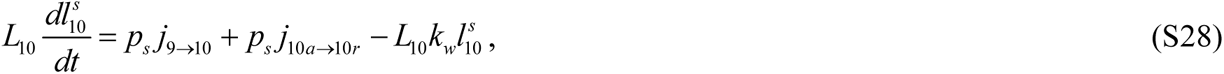

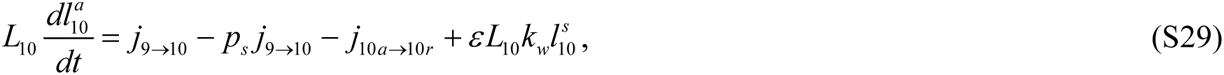

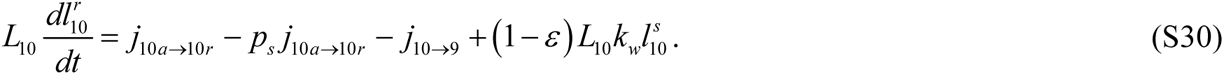

Equations that are analogous to Eqs. (43)-(70) but now for the asymmetric axon displayed in Fig. 1b are as follows. Equations for anterograde fluxes (Fig. 2b) now are

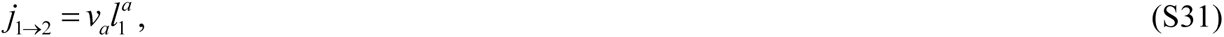

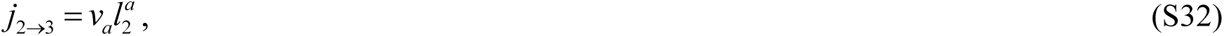

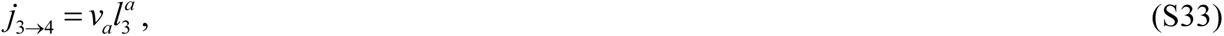

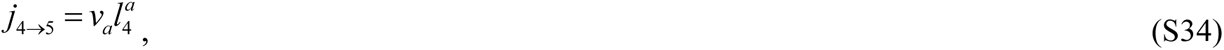

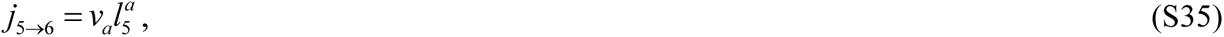

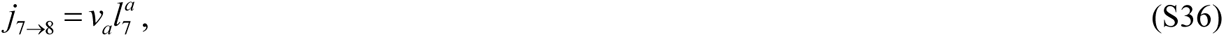

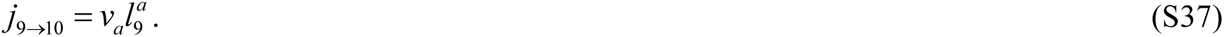

Equations for retrograde fluxes (Fig. 2b) now are

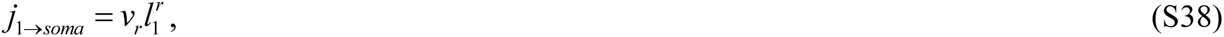

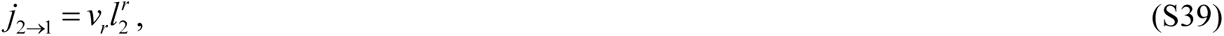

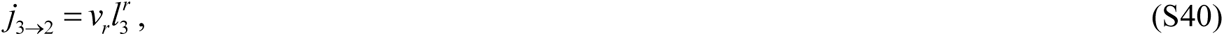

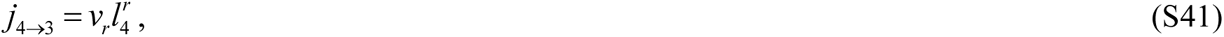

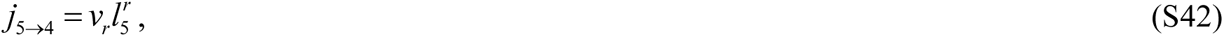

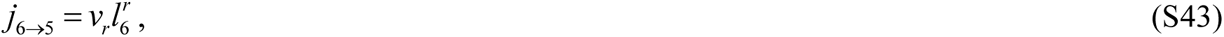

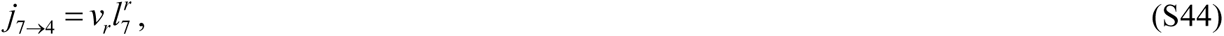

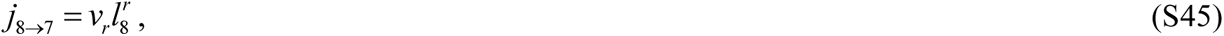

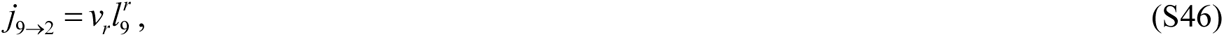

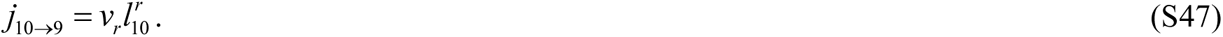

The turn-around fluxes (Fig. 2b) now are simulated as follows:

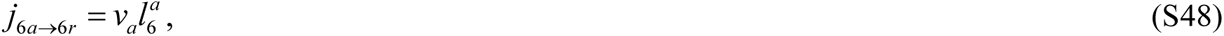

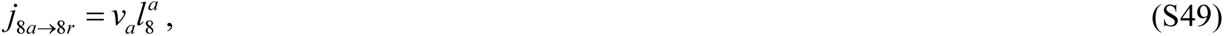

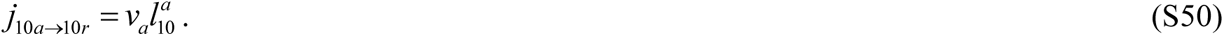

#### S1.2. Model of mean age and age density distributions of mitochondria in the demand sites for the asymmetric axon displayed in Figs. 1b and 2b

Equations that are analogous to Eqs. (74)-(201) but now for the asymmetric axon displayed in Figs. 1b and 2b are (obtained using the method described in Anderson 1983)

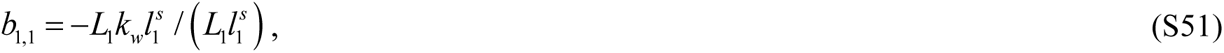

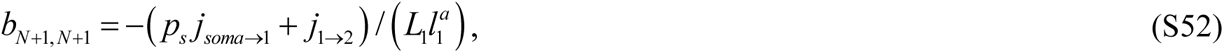

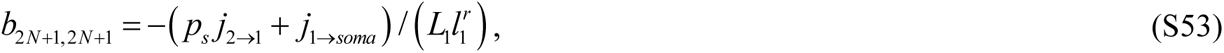

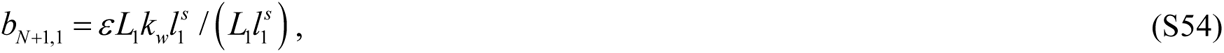

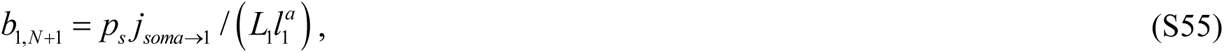

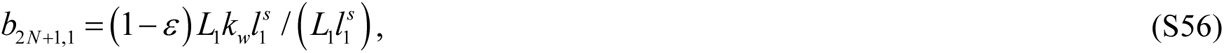

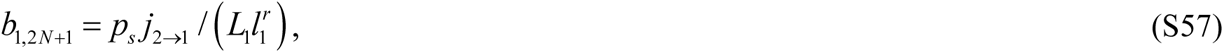

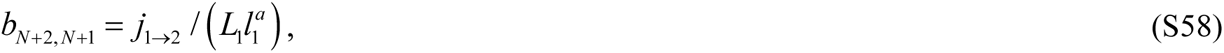

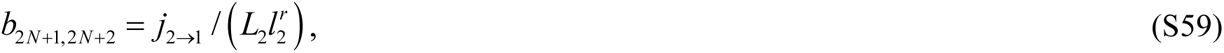

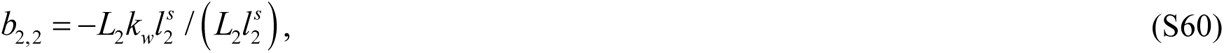

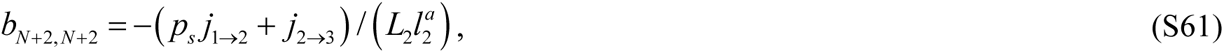

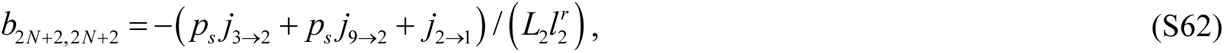

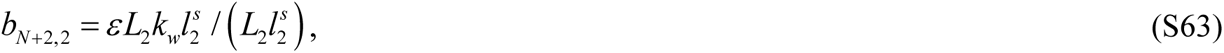

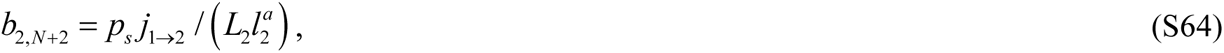

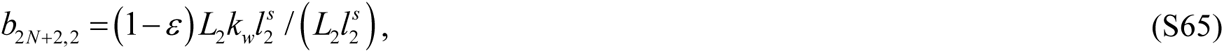

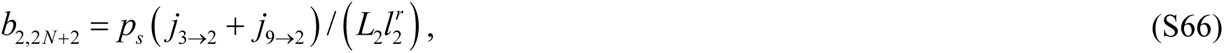

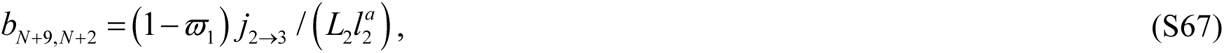

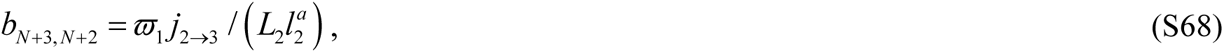

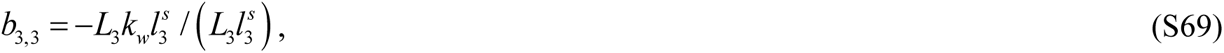

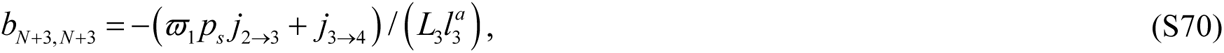

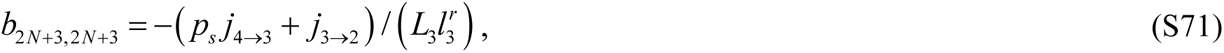

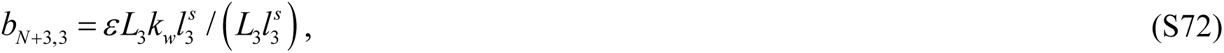

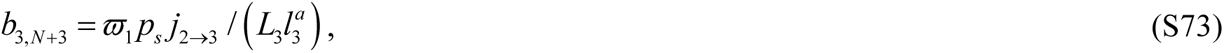

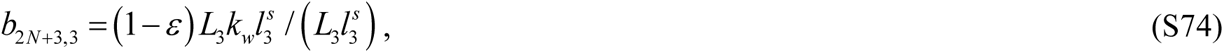

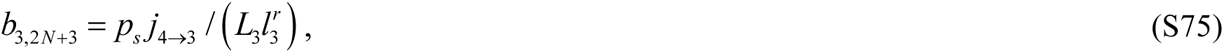

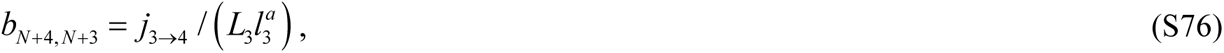

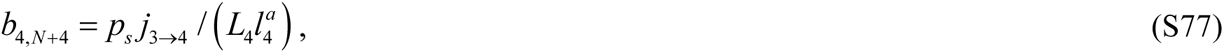

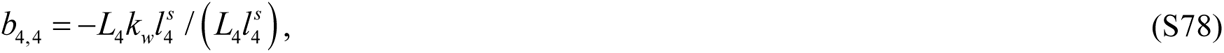

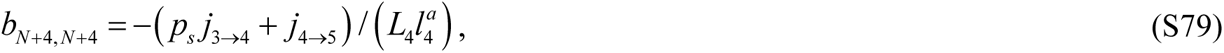

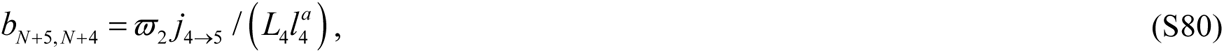

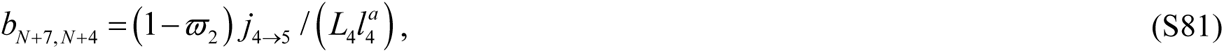

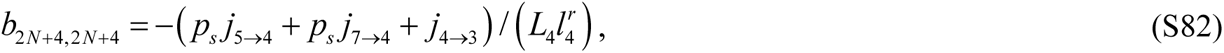

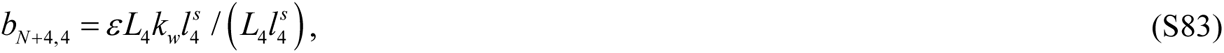

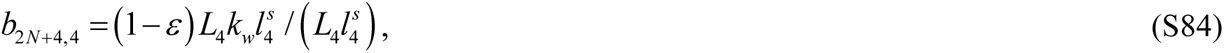

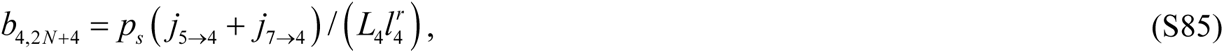

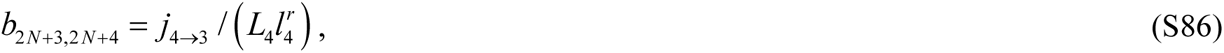

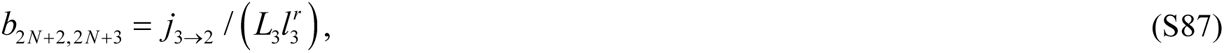

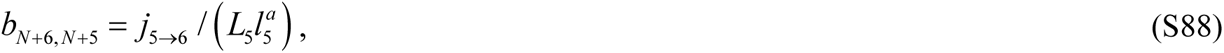

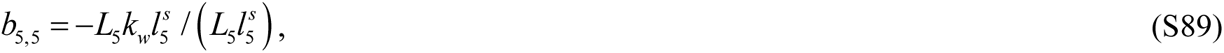

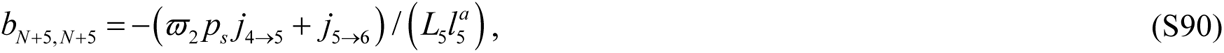

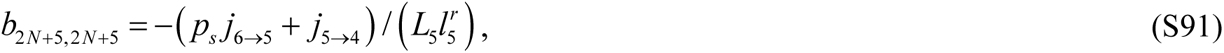

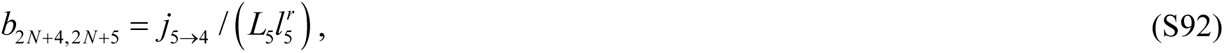

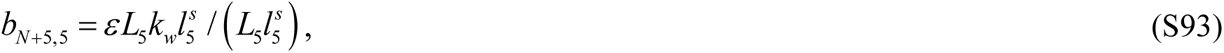

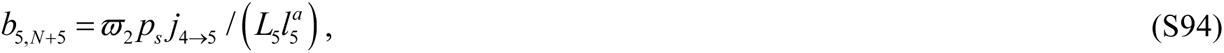

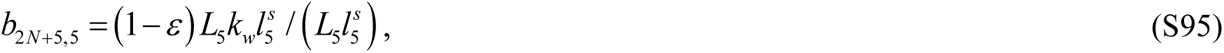

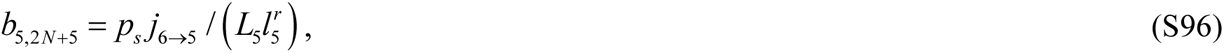

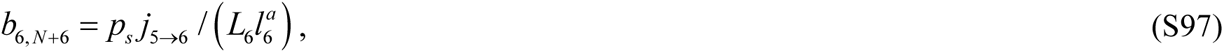

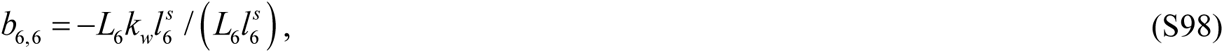

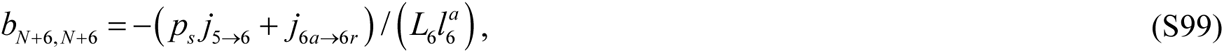

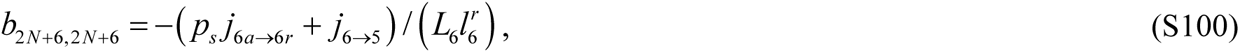

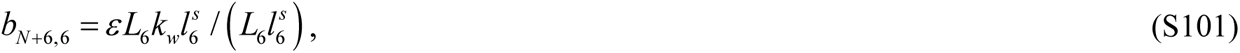

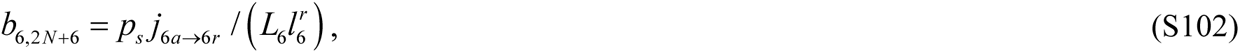

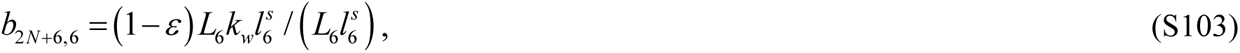

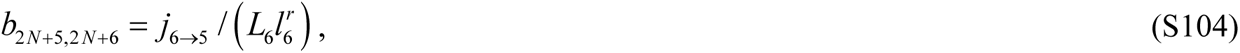

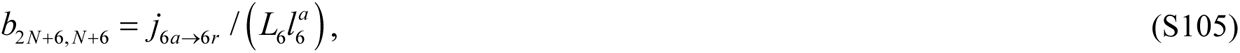

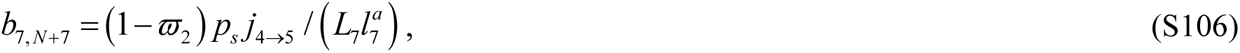

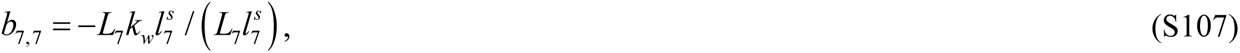

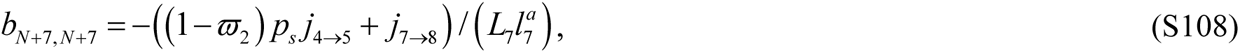

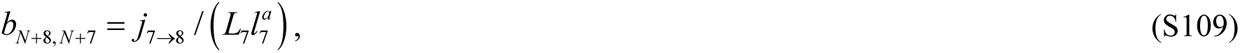

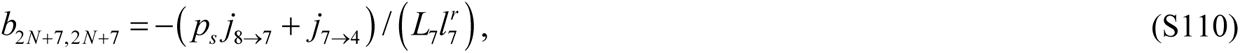

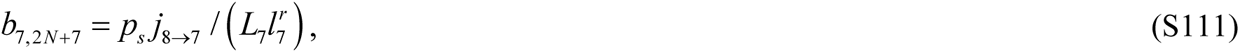

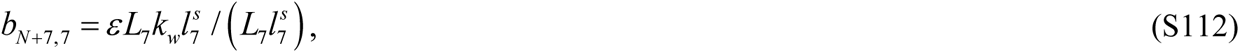

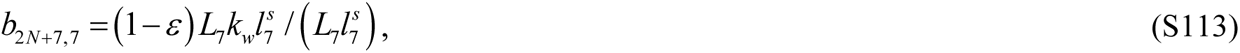

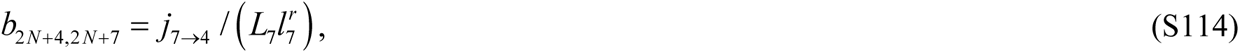

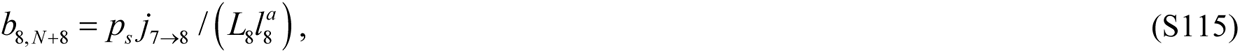

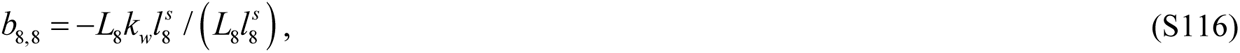

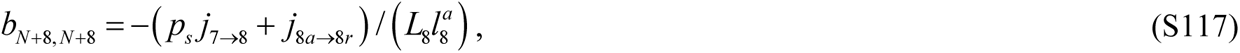

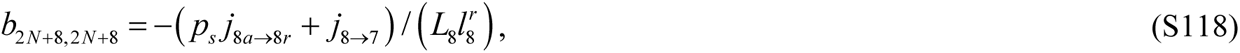

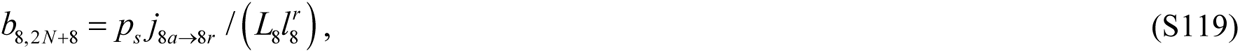

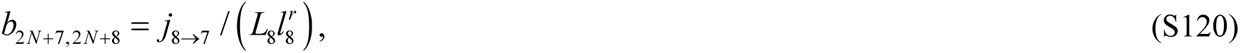

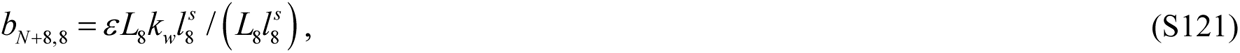

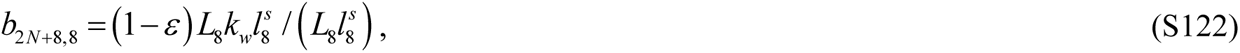

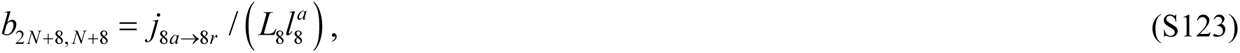

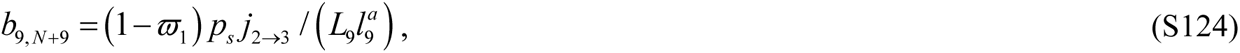

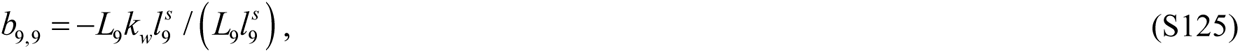

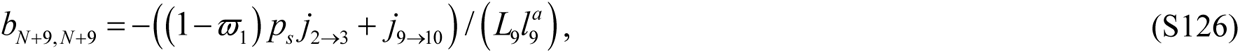

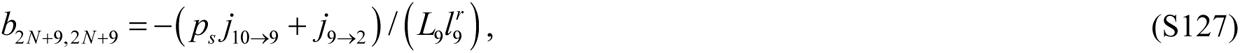

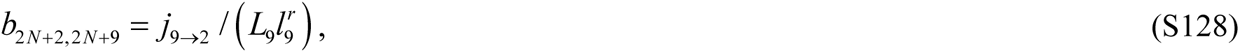

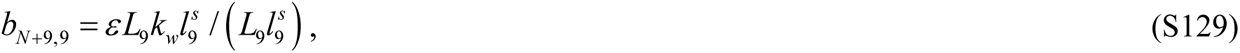

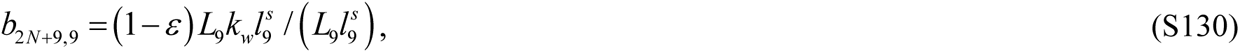

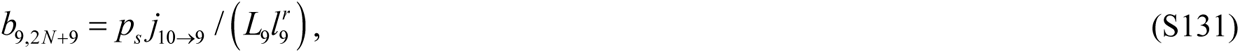

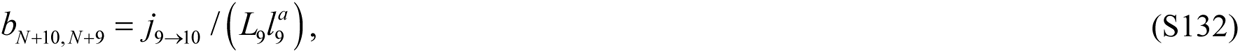

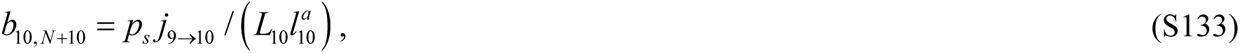

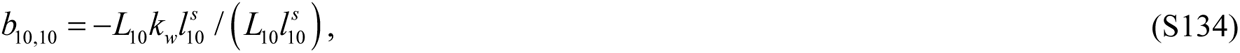

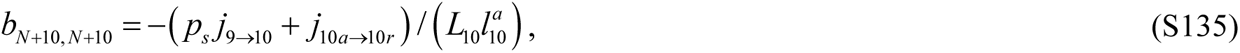

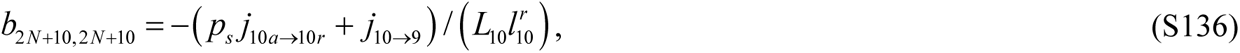

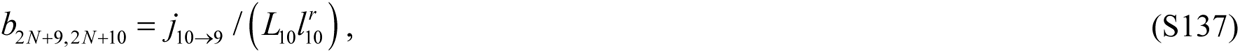

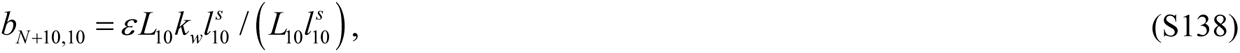

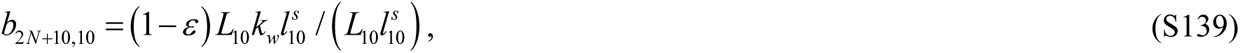

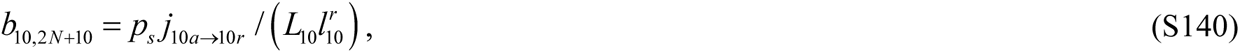

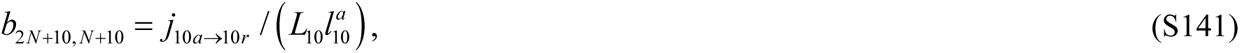

All other matrix B elements, except for those specified in Eqs. (S51)-(S141), are set to zero. The only flux of mitochondria entering the axon is the anterograde flux from the soma to the compartment with anterograde mitochondria by the most proximal demand site, *j_soma_* _→1_.

Our model assumes that all mitochondria that leave the axon (their flux is *j*_1→_*_soma_*) return to the soma for degradation, and none reenter the axon. The mitochondrial age in our model is thus understood as the amount of time that has passed since the mitochondria entered the axon.

The (*N*+1)^th^ element of vector **u** is obtained from the following equation:

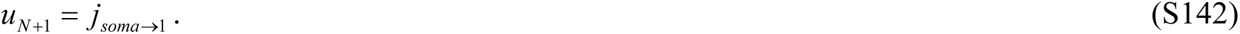

As no other external fluxes enter the terminal,

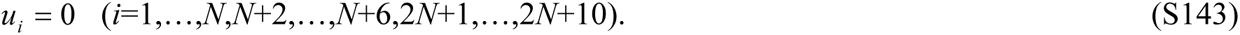

### S2. Supplementary figures

#### S2.1. Symmetric branched axon displayed in **Figs. 1a and 2a**

**Fig. S1.**
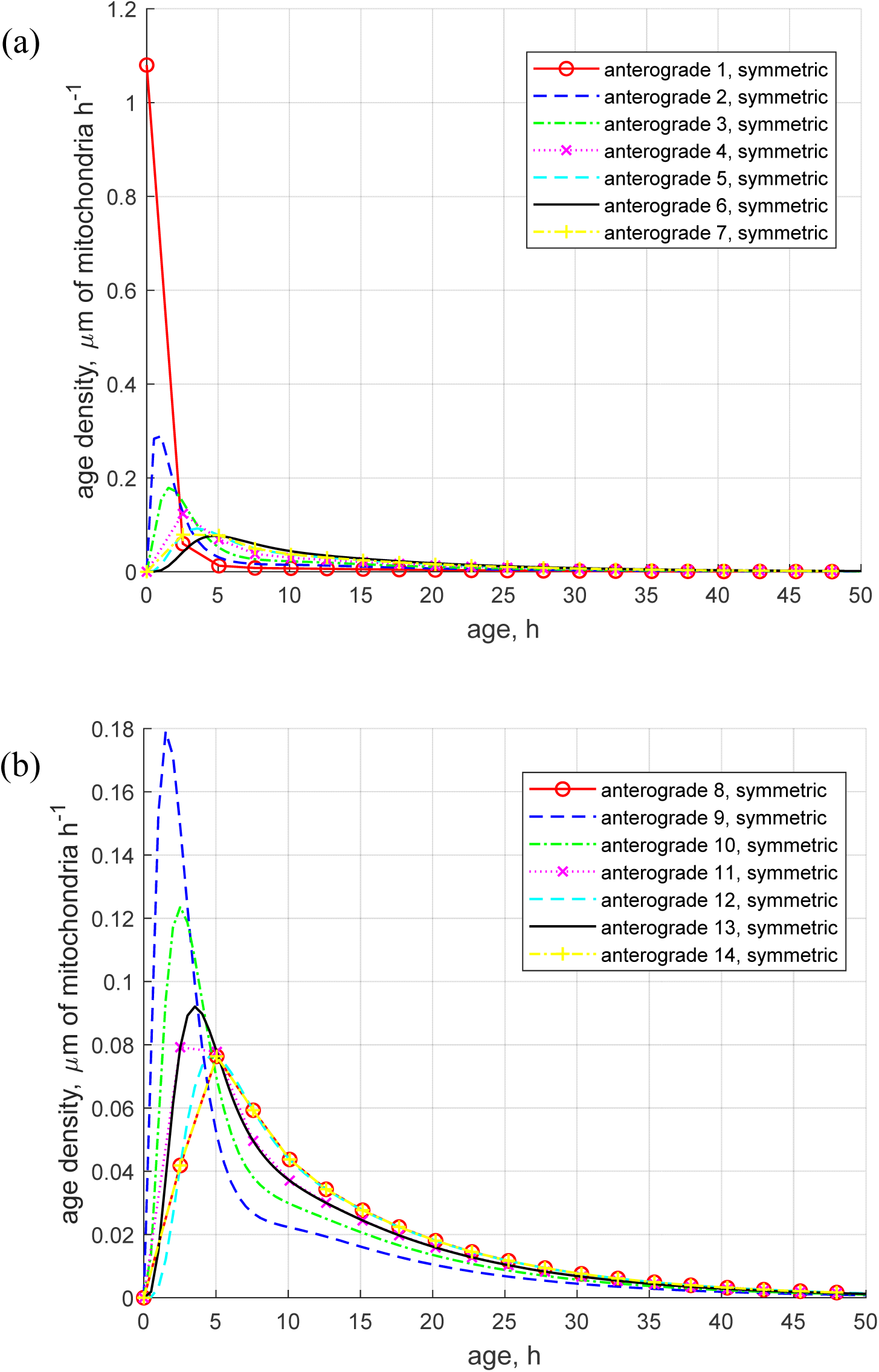
Age density distributions of anterogradely moving mitochondria in a symmetric branched axon across various demand sites. (a) Demand sites 1 to 7; (b) Demand sites 8 to 14. The existence of elongated tails in the positive direction in the age density distributions implies that there are mitochondria in each demand site that are considerably older than the mean age.

**Fig. S2.**
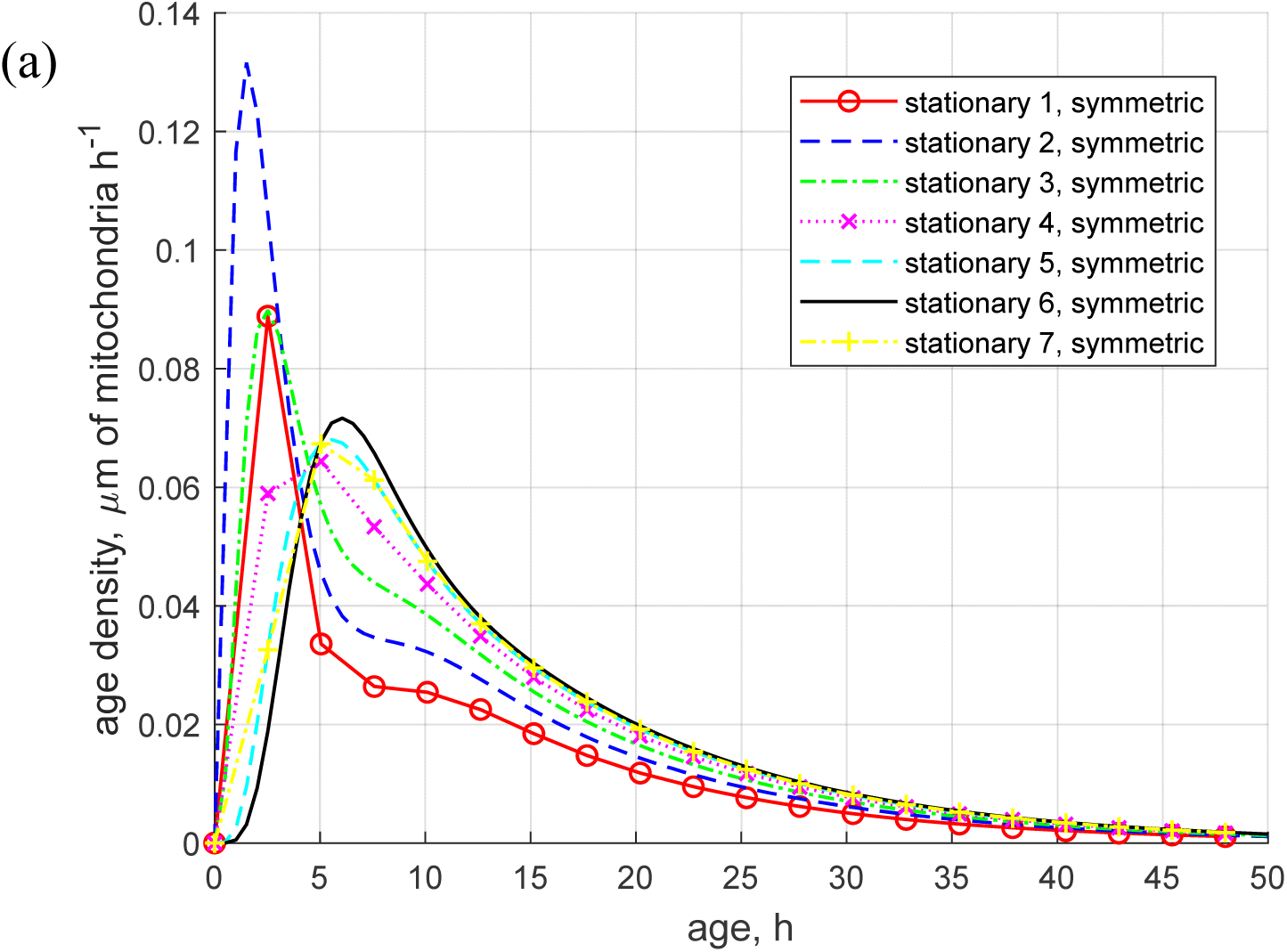
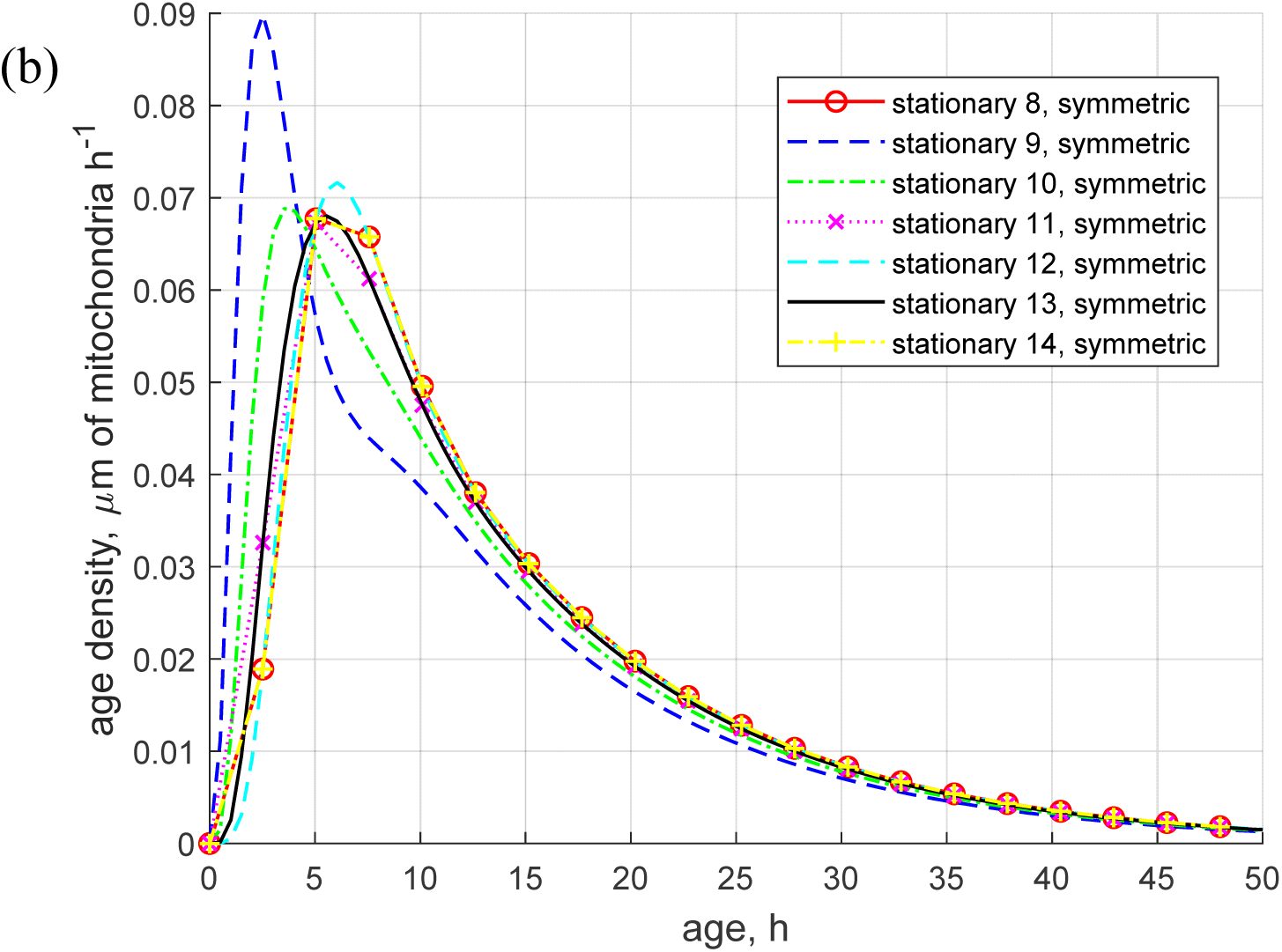
Age density distributions of stationary mitochondria in a symmetric branched axon across various demand sites. (a) Demand sites 1 to 7; (b) Demand sites 8 to 14.

**Fig. S3.**
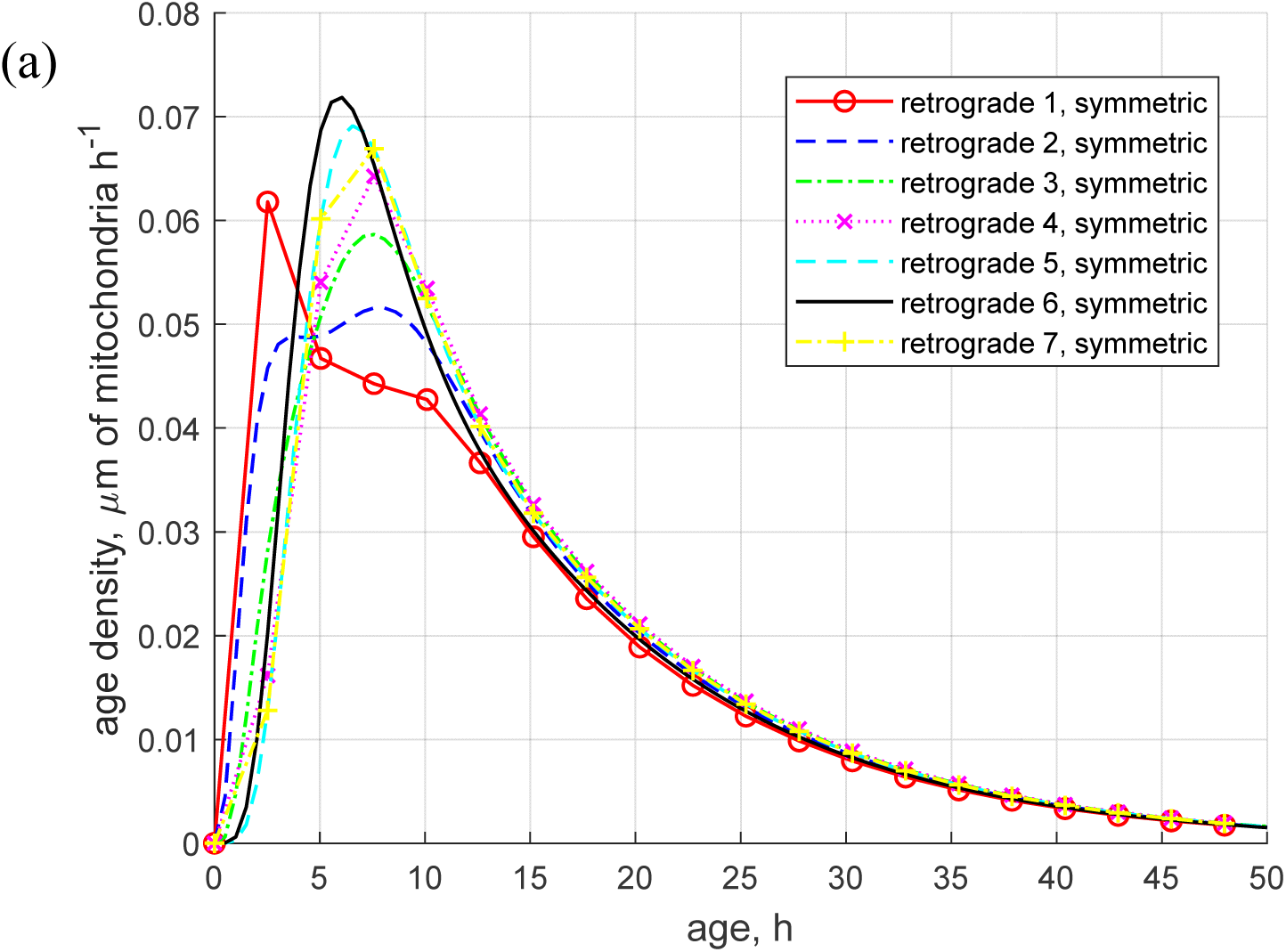
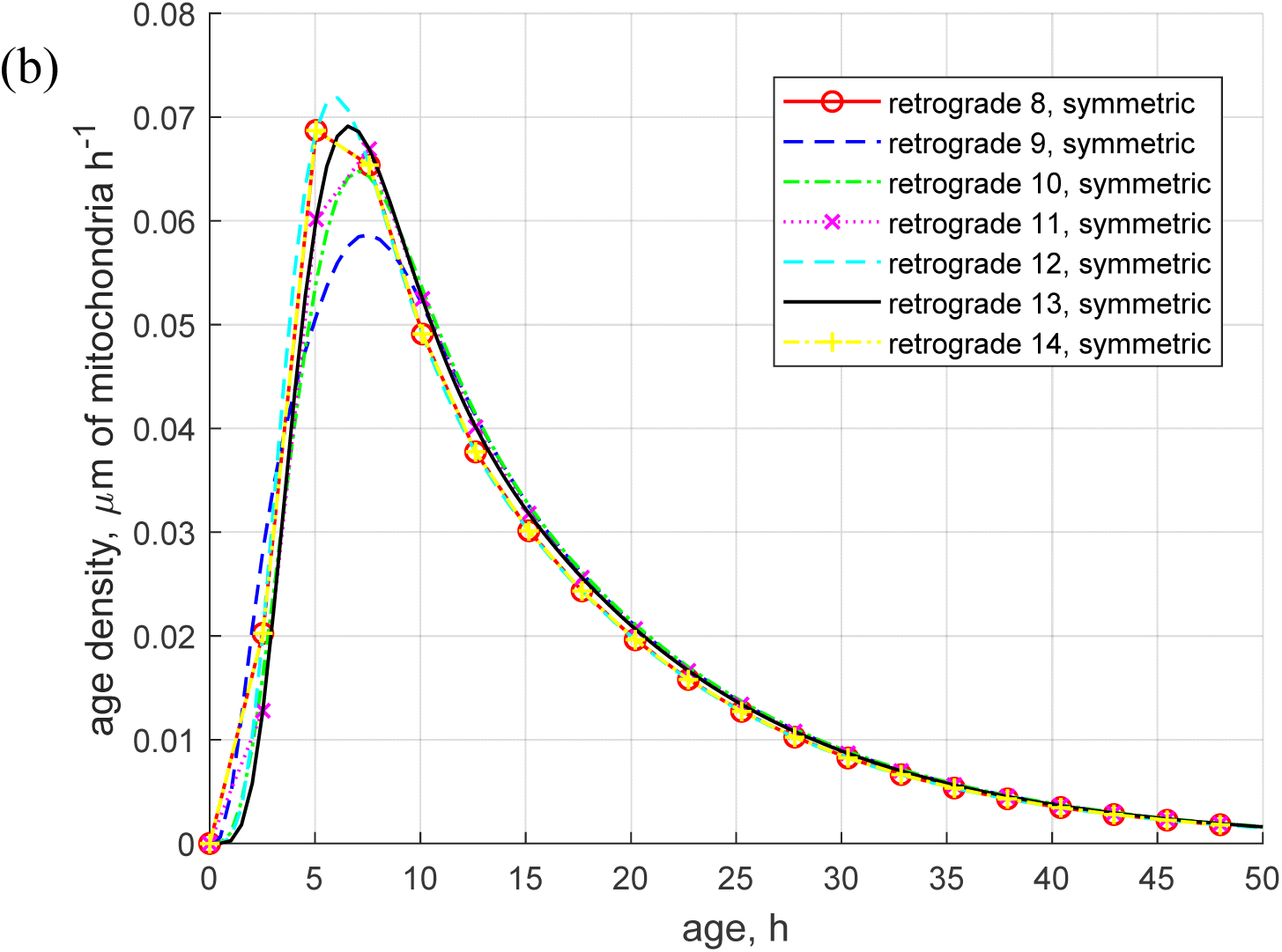
Age density distributions of retrogradely moving mitochondria in a symmetric branched axon across various demand sites. (a) Demand sites 1 to 7; (b) Demand sites 8 to 14.

#### S2.2. Asymmetric branched axon displayed in Figs. 1b and 2b

**Fig. S4.**
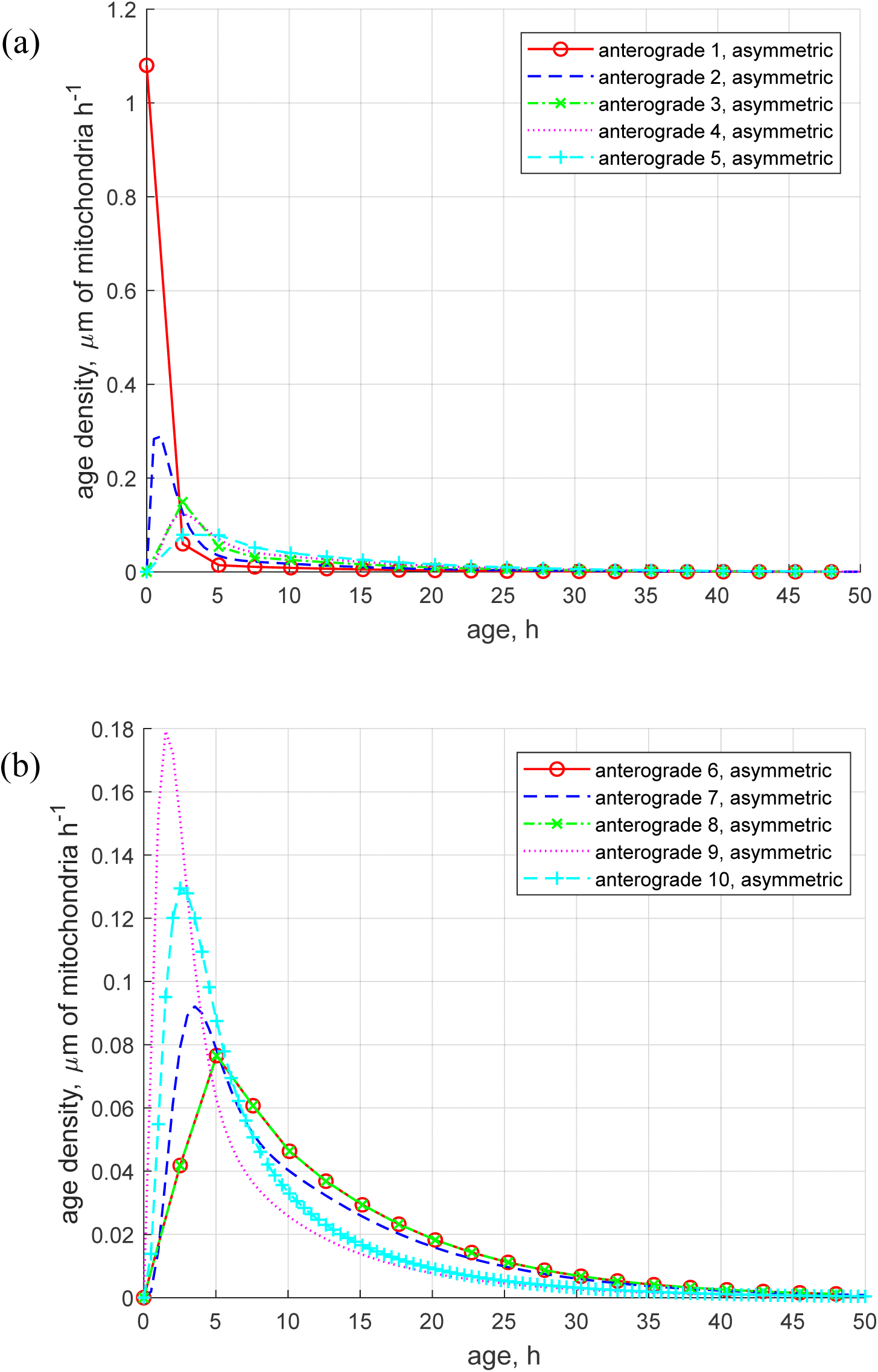
Age density distributions of anterogradely moving mitochondria in an asymmetric branched axon across various demand sites. (a) Demand sites 1 to 5; (b) Demand sites 6 to 10.

**Fig. S5.**
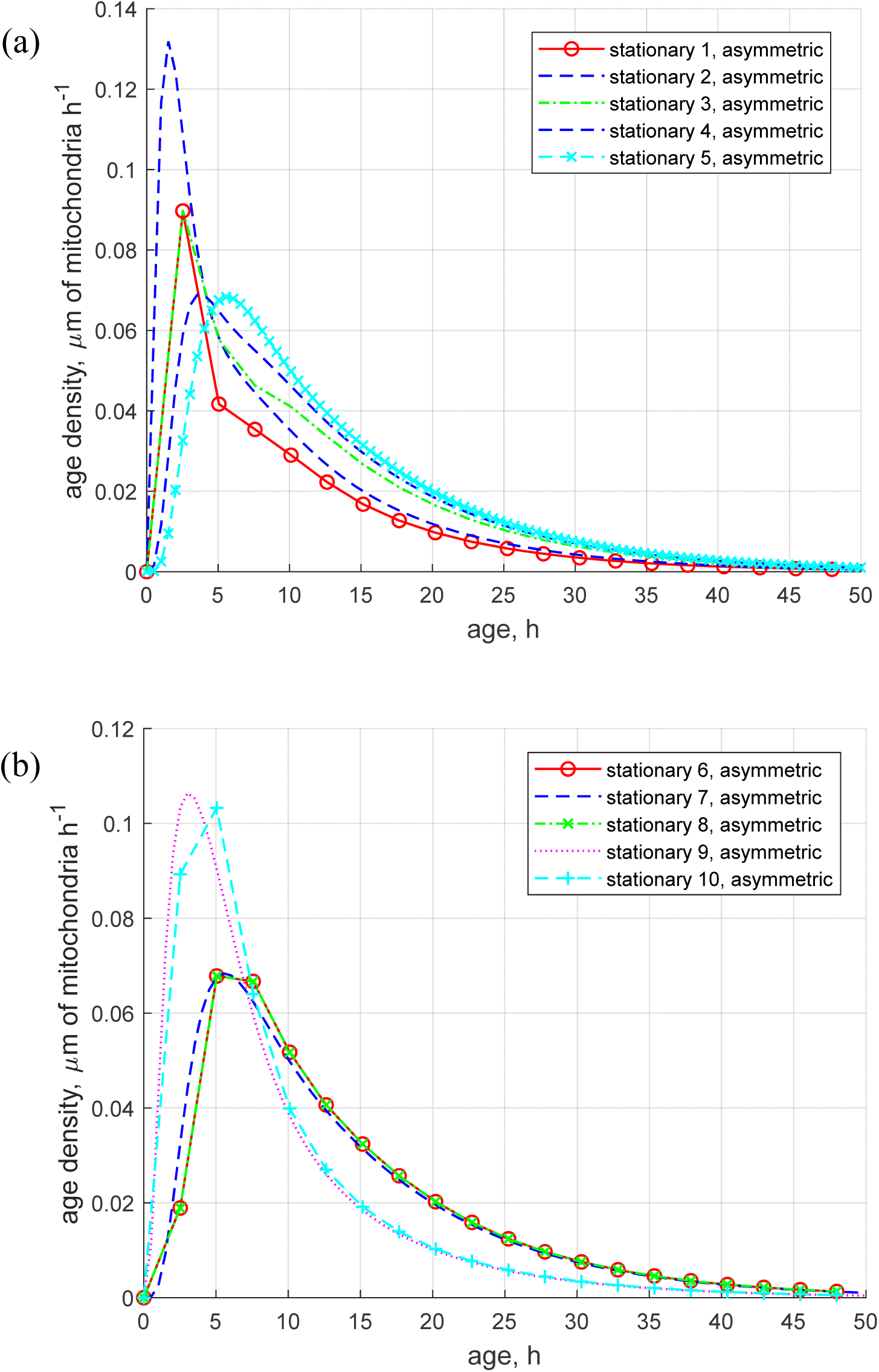
Age density distributions of stationary mitochondria in an asymmetric branched axon across various demand sites. (a) Demand sites 1 to 5; (b) Demand sites 6 to 10.

**Fig. S6.**
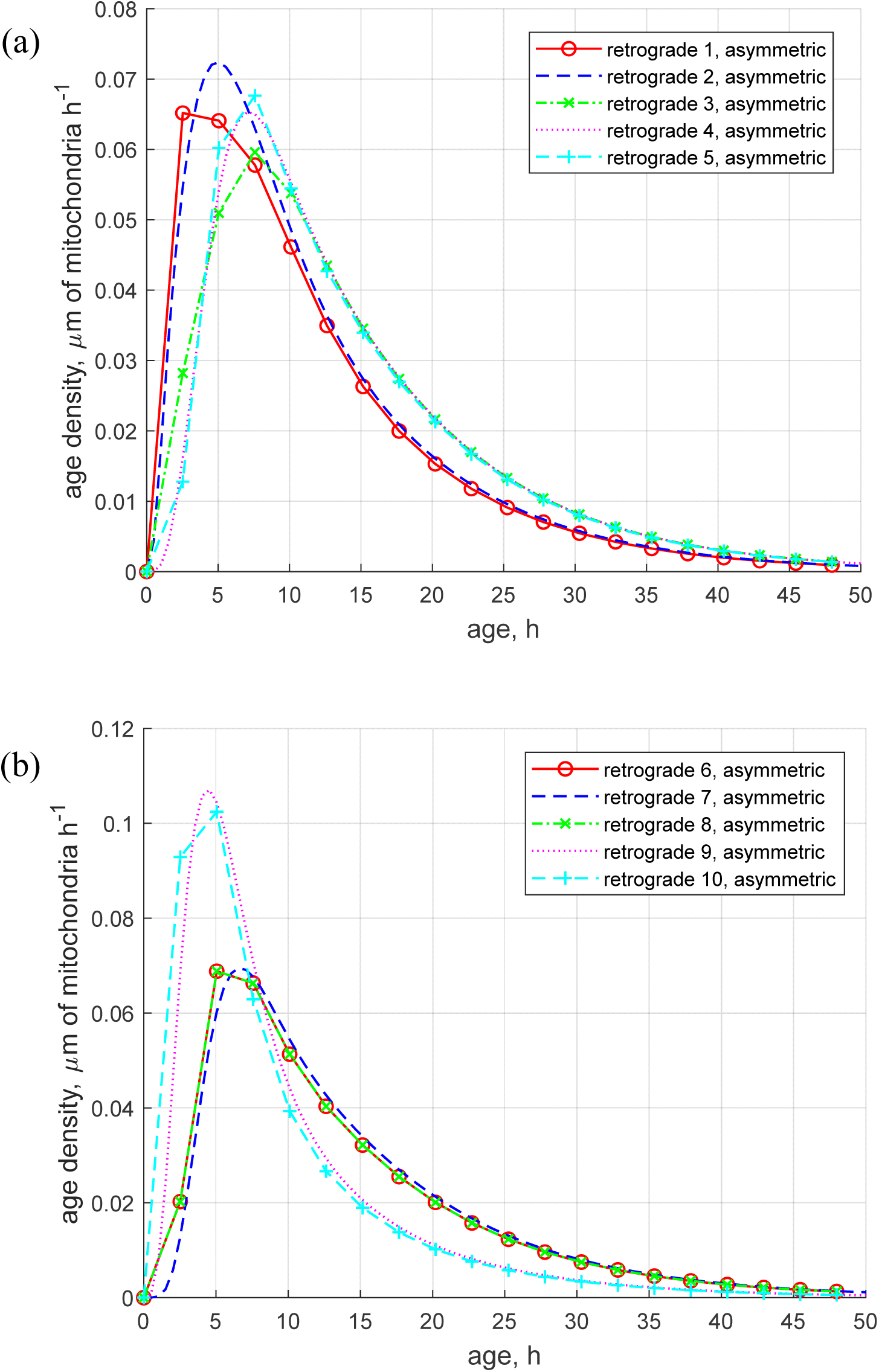
Age density distributions of retrogradely moving mitochondria in an asymmetric branched axon across various demand sites. (a) Demand sites 1 to 5; (b) Demand sites 6 to 10.

**Fig. S7.**
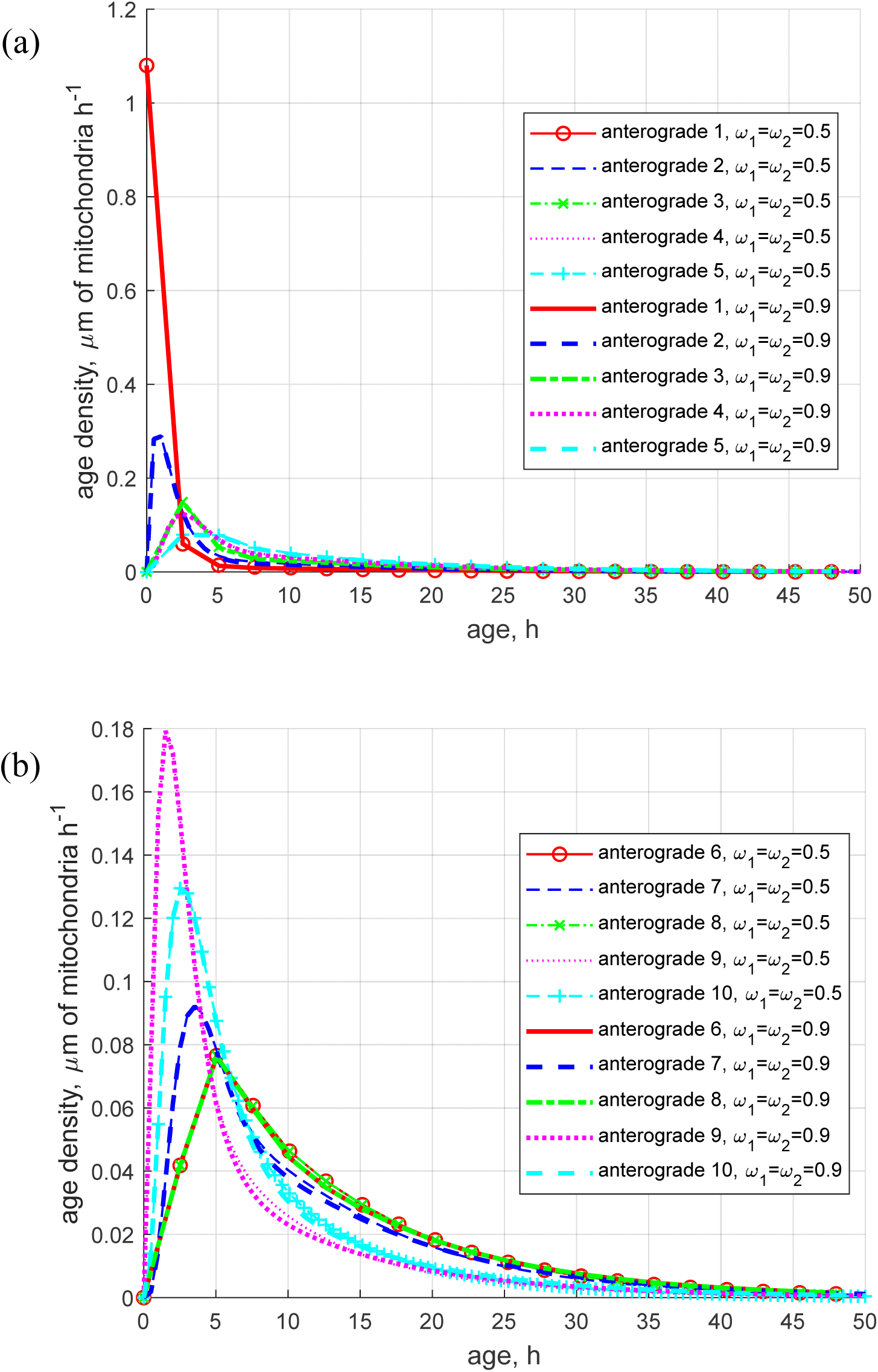
Age density distributions of anterogradely moving mitochondria in an asymmetric branched axon across various demand sites. Comparison of the cases *ω*_1_ = *ω*_2_ = 0.5 and *ω*_1_ = *ω*_2_ = 0.9 for (a) Demand sites 1 to 5; (b) Demand sites 6 to 10. In demand sites 9 and 10, the age density of older mitochondria is visibly higher in the case of *ω*_1_ = *ω*_2_ = 0.9 compared to *ω*_1_ = *ω*_2_ = 0.5. This is because in the case of *ω*_1_ = *ω*_2_ = 0.9, a greater number of mitochondria enter the upper (longer) branch.

**Fig. S8.**
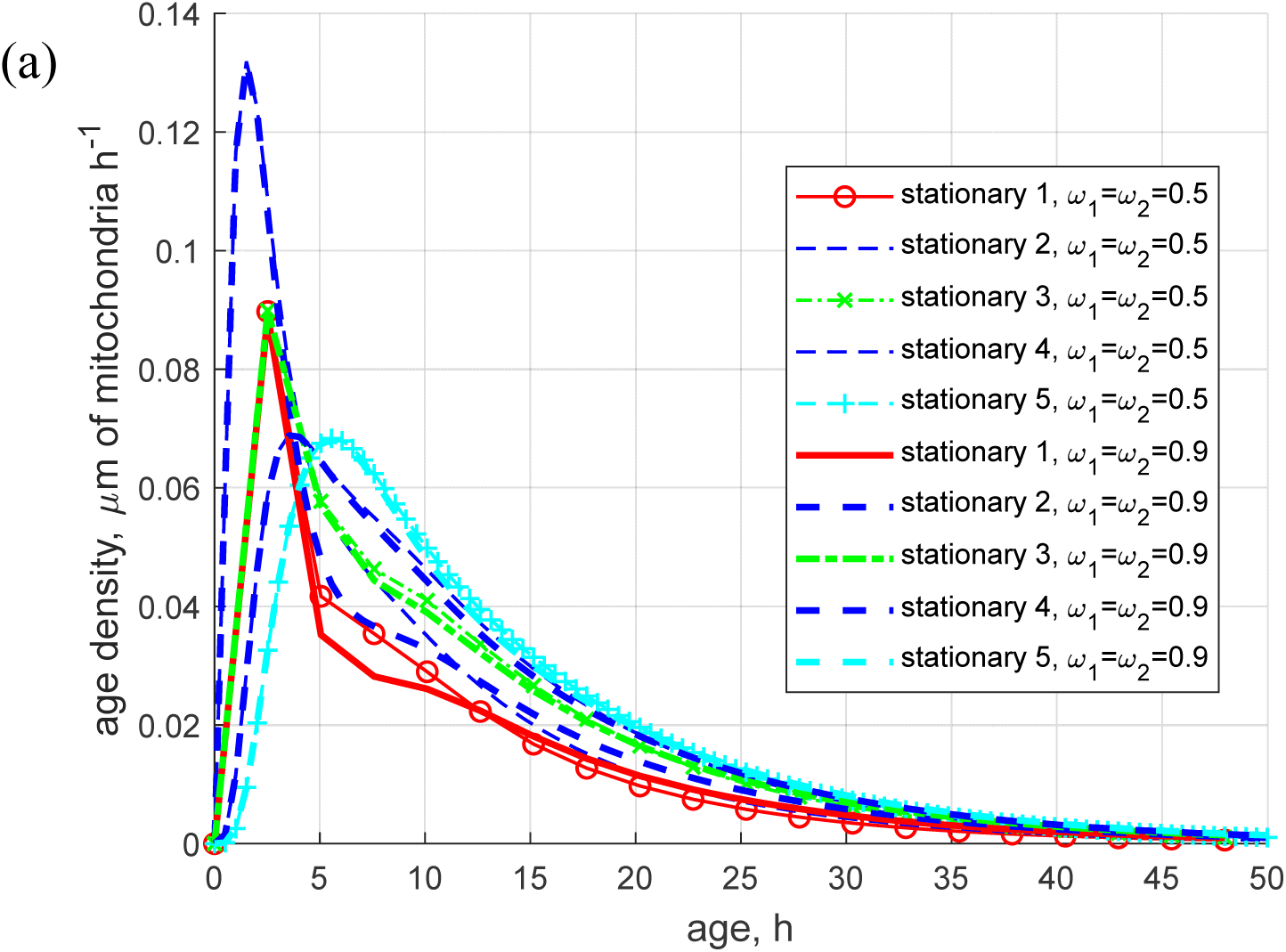
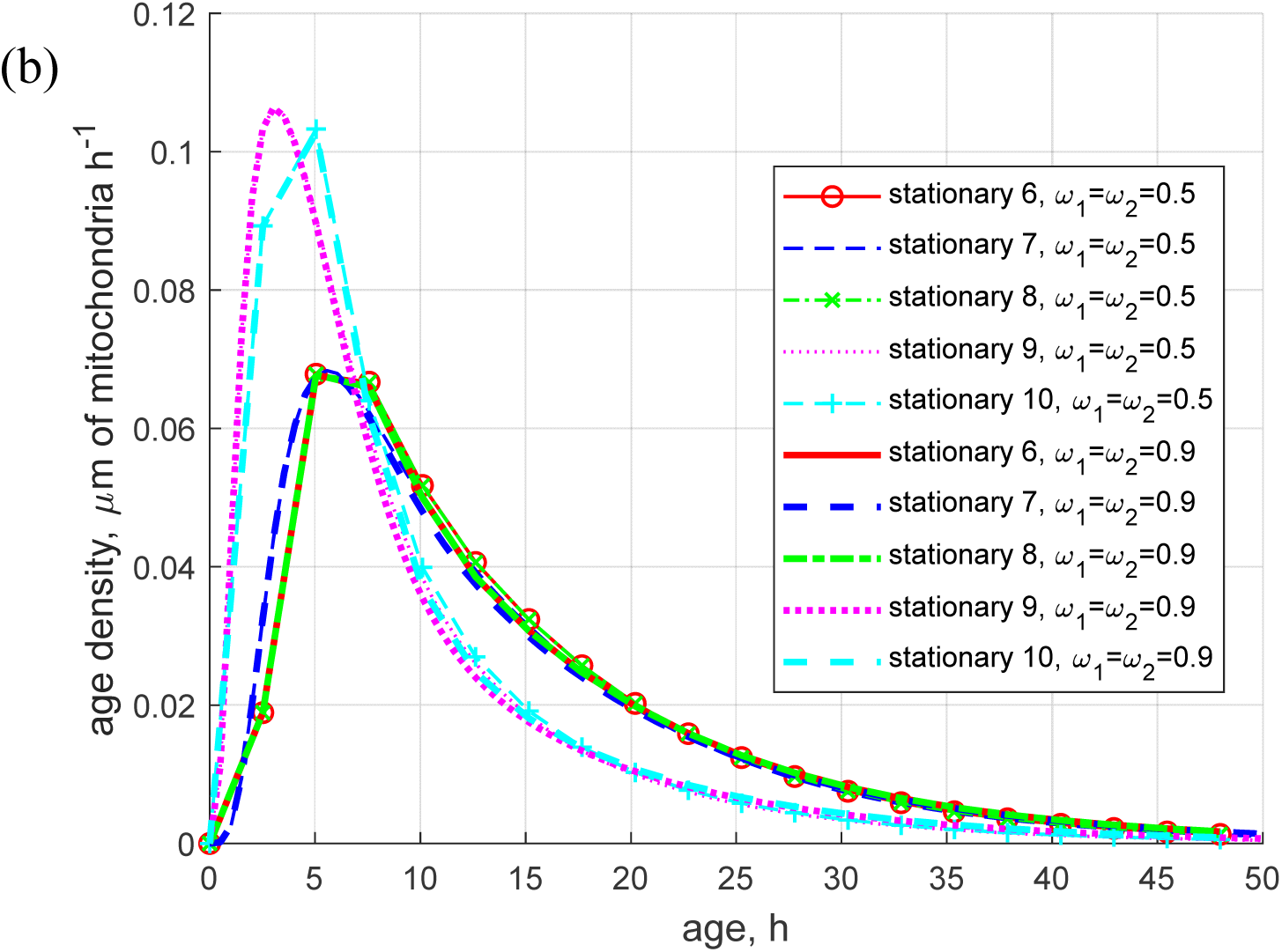
Age density distributions of stationary mitochondria in an asymmetric branched axon across various demand sites. Comparison of the cases *ω*_1_ = *ω*_2_ = 0.5 and *ω*_1_ = *ω*_2_ = 0.9 for (a) Demand sites 1 to 5; (b) Demand sites 6 to 10.

**Fig. S9.**
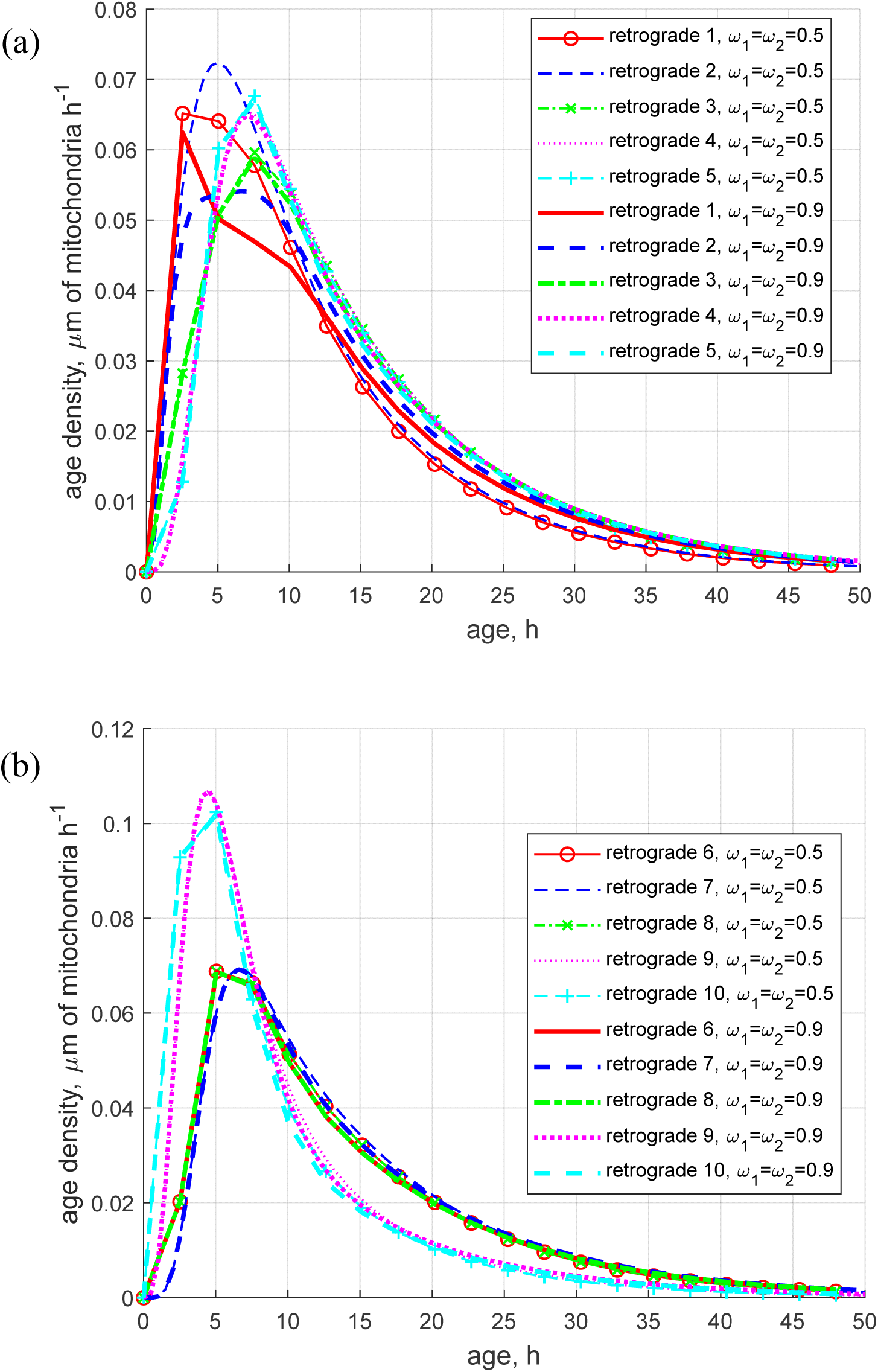
Age density distributions of retrogradely moving mitochondria in an asymmetric branched axon across various demand sites. Comparison of the cases *ω*_1_ = *ω*_2_ = 0.5 and *ω*_1_ = *ω*_2_ = 0.9 for (a) Demand sites 1 to 5; (b) Demand sites 6 to 10.

